# A map of the cortical functional network mediating naturalistic navigation

**DOI:** 10.64898/2025.12.16.694742

**Authors:** Tianjiao Zhang, Emily X. Meschke, Jack L. Gallant

## Abstract

Navigation through the real world requires close coordination of perception, planning, and motor actions. Prior neuroimaging studies that used controlled stimuli and tasks have suggested that navigation-related information is represented broadly across the human cerebral cortex. While three regions in the anterior visual cortex have been well-studied^1^, the extent and functional properties of regions in the parietal and prefrontal cortices are not as well-characterized ^2^. To map and characterize the full cortical network that underpins active navigation in the real world, we used functional magnetic resonance imaging to record brain activity from participants performing a naturalistic navigation task. Banded ridge regression was used to fit high-dimensional encoding models for 28,134 features to this data. Results show that naturalistic navigation is supported by a network of 11 functionally distinct cortical regions: five prefrontal and three parietal regions, along with three regions in the visual cortex that had been identified and characterized in previous studies. Analysis of encoding model weights shows that these 11 regions transform perceptual inputs through decision-making processes to produce action outputs, and are organized along distributed cortical functional gradients. These results provide a unified description of the functional properties and organization of the cortical network that mediates naturalistic navigation. We anticipate that these maps will provide rich targets to inform more targeted future studies of human navigation.

## Main Text

Navigation in the real world is a complex and dynamic task that requires perceiving the world, making plans and decisions, and then acting accordingly ^3^. Understanding the brain systems mediating this continuous perception-cognition-action loop is a fundamental goal in neuroscience. Studies have shown that spatial navigation recruits a network of regions in the human brain, including the hippocampus ^4,5^, entorhinal cortex ^6,7^ retrosplenial cortex (RSC, ^8,9^), occipital place area (OPA, ^10^), and parahippocampal place area (PPA, ^11,12^). While there is evidence that parietal and prefrontal regions are also involved in navigation ^2,13^, these regions have not been delineated precisely, and their precise role in navigation are not well characterized. Thus, to understand how the human brain navigates through the world, there is a critical need to fully map the cortical extent of the functional brain network mediating spatial navigation.

Fully understanding the navigation network in the human brain also requires answering another fundamental question: how is the representation of navigation-related information organized across its constituent regions? A modular perspective suggests that each aspect of navigation is mediated by a distinct functional region ^14–17^. Much of the existing evidence supports this view: for example, the identify of a visual scene is thought to be represented by the PPA ^18,19^, navigation affordances are thought to be found in the OPA ^20^, and abstract cognitive map representations such as the distance to a goal are found in hippocampus ^21^. Yet, more recent evidence points towards a distributed network-based organization ^3,15^. Neuroimaging studies have found that both PPA and OPA are situated within larger identity- and affordance-selective networks ^19^, and head direction representations are found across multiple functional regions ^22,23^. Likewise, neurophysiology studies in rodents have found that navigation-related information is represented in overlapping gradients that extend across ROI boundaries ^24,25^. Resolving these competing hypotheses of organization requires fully characterizing the representation of navigation-related information across the network.

A significant barrier to fully characterizing the navigation network in the human brain is that most studies have used tightly controlled tasks and stimuli to target individual regions or specific representations. However, because the brain is a nonlinear system, it behaves differently under simplified conditions than in more complex naturalistic conditions ^26–28^. Indeed, rodent studies have shown that landmark tuning in RSC neurons degrades when animals are not actively navigating ^29^, and complex navigation tasks recruit much more of the dorsal cortex than simpler tasks ^30^. Consequently, controlled tasks may not engage the full navigation network in the human brain, and the lack of emergent interactions between network components in controlled experiments may artificially create the appearance of distinct modules. Overcoming this barrier thus requires naturalistic experiments that engage the brain under ecologically valid conditions ^27,31^.

### Naturalistic navigation during fMRI

To fully map the human cortical network that underlies active navigation in naturalistic environments and characterize its functional organization, we developed a naturalistic driving simulation system for functional magnetic resonance imaging (fMRI). We used Unreal Engine 4 (Epic Games) and the Carla plugin ^32^ to build a 2×3 km virtual world (Fig. 1A, Supplementary figure 1). The world contains a city comprising several distinct urban and suburban neighbourhoods, along with more rural areas (Fig. 1B). The world is populated by AI-controlled vehicles and pedestrians who obey traffic rules. Prior to scanning, the simulator was distributed to each participant, and they used it to learn the world by driving through it for approximately 10 hours. To help participants learn the map, navigational aids were displayed on screen, and the simulator directed them to visit every possible destination. (No navigation aid was displayed during the actual fMRI experiment.) In the scanner, participants used a custom MR-compatible steering wheel and pedal set to drive from a first-person perspective (Fig. 1C). Functional magnetic resonance imaging (fMRI) was used to record blood oxygen level-dependent (BOLD, ^33^) activity from participants while they performed a taxi-driver task ^34^ in the virtual world (Fig. 1D). Eyetracking data, behavioural actions, and ground-truth world state were also recorded continuously.

**Fig. 1.**
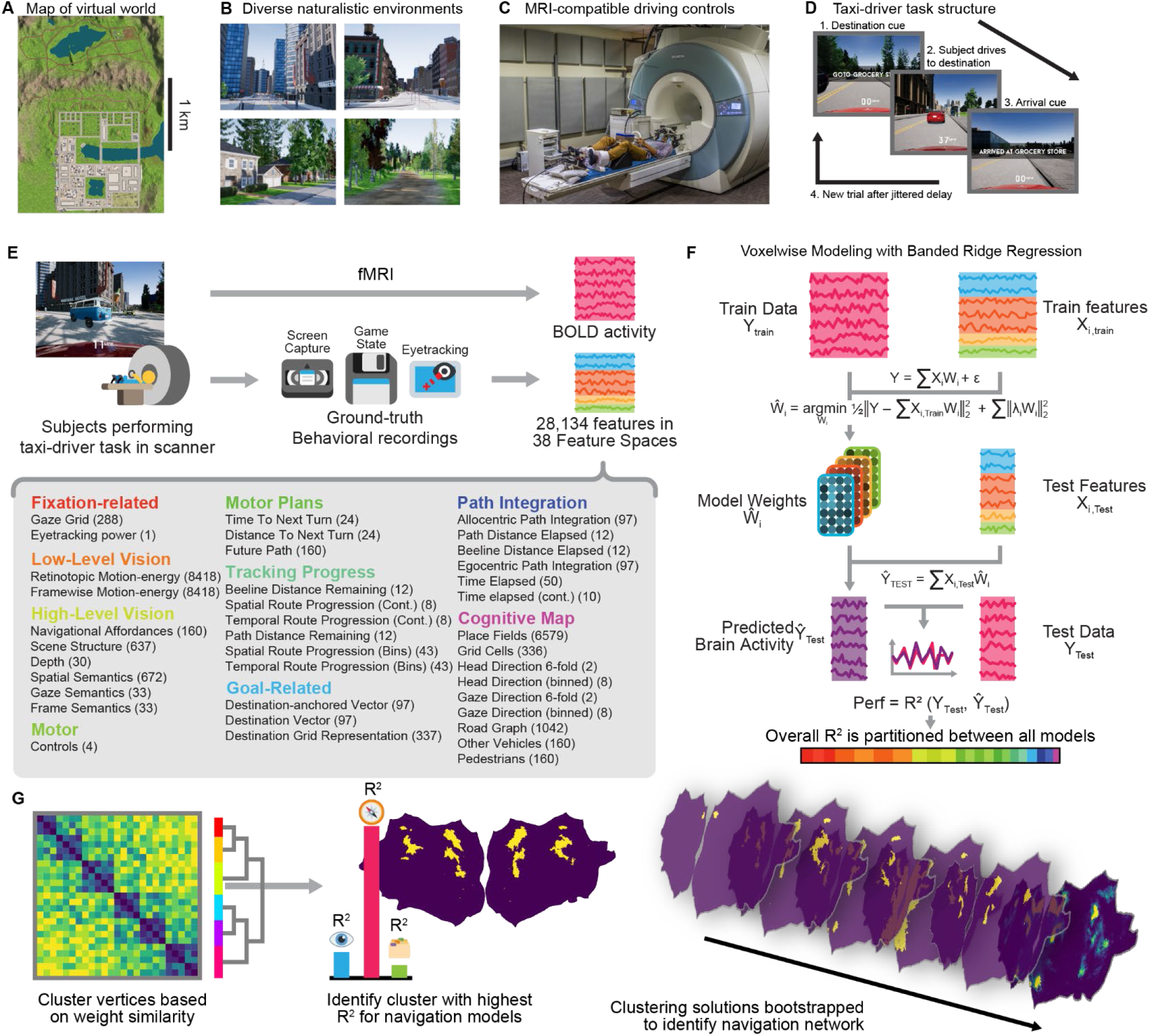
Using fMRI to map the cortical networks mediating active, naturalistic navigation in a virtual world. (**A**) We built a large virtual world (approx. 2 × 3 km) that contains over 200 uniquely identifiable locations and is populated by AI vehicles and pedestrians. (**B**) The virtual world contains diverse environments, including commercial, urban, suburban, and rural areas. (**C**) Participants used a set of custom MR-compatible steering wheel and pedals to drive a virtual car. Participants learned the layout of the world prior to scanning by driving through the world. (**D**) In the scanner, participants performed a taxi-driver task while BOLD activity was recorded with fMRI. On each trial, participants were asked to drive to a randomly selected destination via the fastest path while obeying all traffic rules. A new trial began after the participant arrived at the destination. (**E**) Eyetracking data, screen capture, and ground-truth world information from the game engine were recorded in addition to BOLD activity. These recordings were used to extract 28,134 time-varying features across 38 feature spaces that described many aspects of the experiment, such as motion-energy features that capture the low-level visual structure of the visual scene, route progression features that describe the progress of the participant to the current destination, or motor features that describe the participant’s actuation of the steering wheel and pedals. (**F**) Banded ridge regression was then used to fit voxelwise encoding models for all feature spaces simultaneously to the BOLD activity in every voxel. Model performance was then quantified by prediction accuracy on a hold-out dataset that was not used during model fitting. The overall model performance was then partitioned across models for each of the 38 feature spaces. (**G**) To identify functional networks that specifically represent navigation-related information, model connectivity was applied to the encoding model weights. Weights from all participants were averaged on the fsaverage6 surface and then hierarchically clustered. Because clustering solutions are sensitive to both noise and the clustering parameters, the clustering solution was bootstrapped. For each vertex, a *navigation preference index* (NPI) was defined as the frequency that it was grouped into the cluster with the highest encoding model performance aggregated across all navigation feature spaces. The navigation network was then identified as the set of vertices with high NPIs.

The taxi-driver task required participants to plan and follow routes purely from memory, without any navigational aids. To verify that the participants were able to navigate successfully, we quantified their behavioural performance. A path optimality value was defined for each trial as the ratio between the length of the actual path taken by the participant and the length of a path found by A* search ^35^. A path optimality of 1 indicates that the participant found an optimal path, while larger values indicate suboptimal paths. Because our implementation of the A* algorithm could not account for all environmental constraints, some paths appeared to be super- or sub-optimal (see Supplementary figure 2 for details).The mean path optimality was 1.06 ± 0.07 (mean ± std across participants), suggesting that participants were near-optimal in pathfinding (p = 0.12, two-sided t-test, n= 6, Supplementary figure 2A).

### Identifying cortical navigation ROIs

To model the BOLD signals recorded during the taxi-driver task, we defined 38 separate feature spaces that quantify perceptual, cognitive, and motor information that prior human and animal studies suggest might be represented in the brain during active navigation (Fig. 1E, Supplementary table 1, Supplementary Methods). Taken together, these feature spaces comprise 28,134 separate features. For expository purposes, we group the feature spaces into nine categories: those related to fixation, low-level vision, high-level vision, motor responses, motor plans, route progression, navigational goals, path integration, and cognitive maps. (A short description of each feature space is found in Supplementary table 1. For a detailed description of each feature space, see Supplementary Methods.) Banded ridge regression ^36,37^ was used to simultaneously fit encoding models for all 38 feature spaces (Fig. 1F). A separate encoding model was fit to each voxel and to every individual participant. The fit model weights describe how each feature affects BOLD signals recorded in each voxel. Model prediction performance is quantified by the variance explained (R^2^) on a hold-out test data set. To recover the functional network mediating active navigation, bootstrapped model connectivity ^38^ was used to functionally cluster the cortex and identify the network of regions that represented navigation-specific information (Fig. 1G). Encoding model performance, partitioned across models for the 38 feature spaces, was then used to interpret the functional properties of the cortical regions that comprise the navigation network.

To determine whether the taxi-driver task fully elicited activity broadly across the cerebral cortex, we first evaluated the performance of the joint model that aggregated models for all 38 feature spaces. Inspection of Figure 2A reveals that the joint model explains brain activity in many regions of the cerebral cortex (for individual participants see Supplementary figure 3; for split R^2^ scores for each of the model categories, see Supplementary figure 4). In the visual cortex, the join model explains activity in multiple areas that represent perceptual information, such as V1, human middle temporal complex (hMT), intraparietal sulcus (IPS), and the frontal eye fields (FEF), and also three known navigation-related functional areas: RSC, OPA, and PPA. In the parietal cortex and prefrontal cortex, the joint model explains activity in areas that are thought to represent information related to memory ^39^, planning ^40,41^, or executive processes ^42^, such as the precuneus, medial prefrontal cortex (mPFC), the inferior frontal sulcus (IFS), and the anterior insula. The joint model also explains activity in regions known to represent motor-related information, such as the somatomotor cortex and the premotor cortex. Thus, our taxi-driver task indeed elicits activity broadly across the cerebral cortex, including multiple sensory, cognitive, and motor regions.

**Fig. 2.**
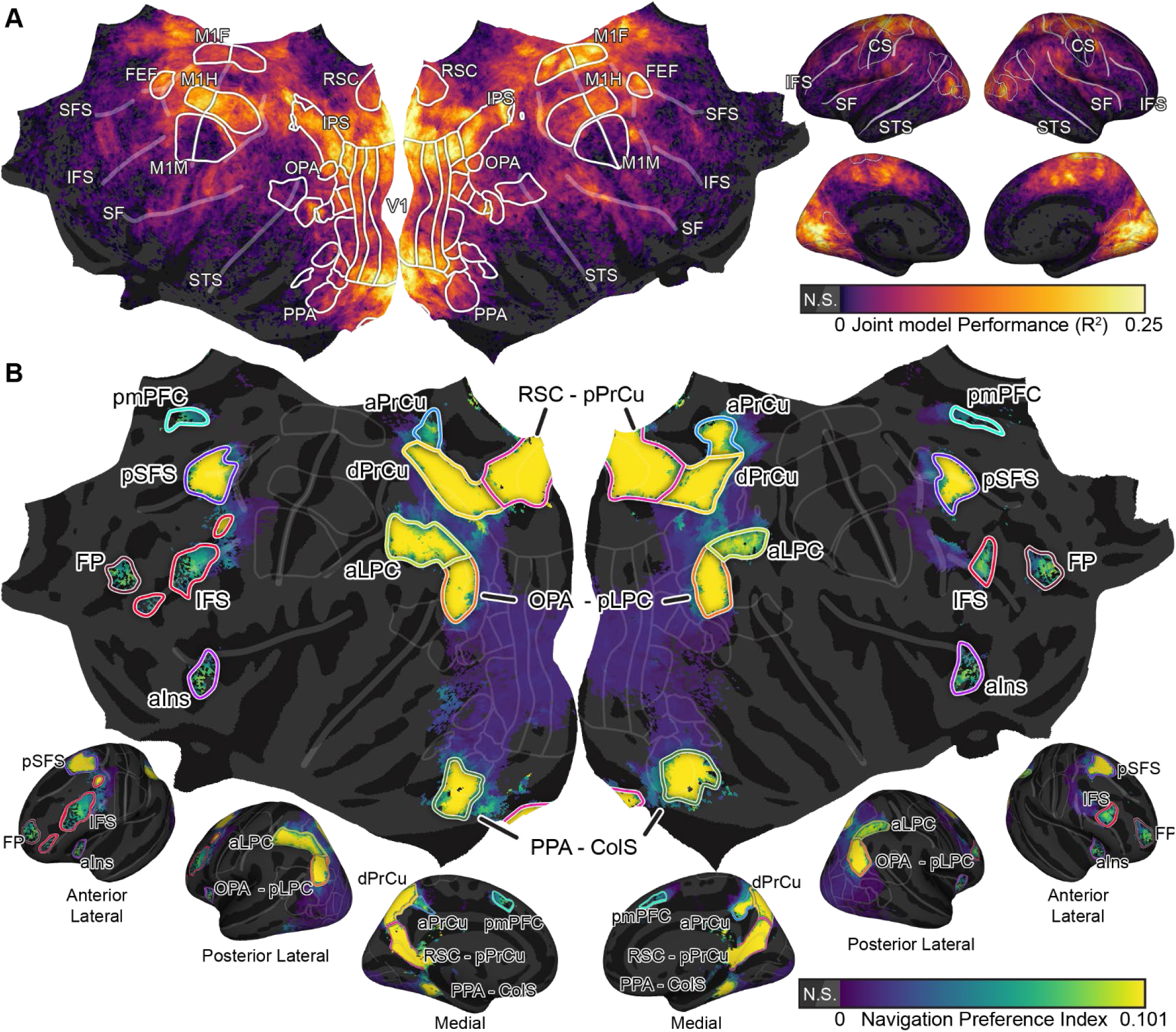
Active, naturalistic navigation recruits a distributed network of brain regions. **(A**) Active, naturalistic navigation engages much of the cortex. The joint model prediction performance for all 38 models is shown on the flattened and inflated views of the cortical surface. Models explain significant activity across the visual, parietal, and prefrontal cortices (Benjamini-Hochberg corrected p < 0.05, permutation test, curvature indicates non-significant vertices). These include many regions that encode non-navigation-specific information, such as visual regions, the motor cortex, and parts of the default mode network. (**B**) To identify the network of cortical regions that specifically supported active, naturalistic navigation, we used encoding model prediction performances and model connectivity to calculate a navigation preference index (NPI) for each vertex. The NPI is shown on the flattened and inflated views of the cortical surface. High NPI vertices are distributed across the occipital, parietal, and prefrontal cortices (Benjamini-Hochberg corrected p < 0.05, permutation test, curvature indicates non-significant vertices). This map was used to define the ROIs that comprise the cortical network that mediates active, naturalistic navigation. Spatial variations in the NPI were used to delineate boundaries between ROIs. In total, 11 distinct ROIs were identified, the most posterior of which overlap significantly with known scene-selective ROIs. For ease of reference, we operationally refer to these ROIs by their anatomical locations and overlaps with known scene-selection ROIs: PPA-collateral sulcus (PPA-ColS), OPA-posterior lateral parietal cortex (OPA-pLPC), RSC-posterior precuneus (RSC-pPrCu), dorsal precuneus (dPrCu), anterior precuneus (aPrCu), anterior lateral parietal cortex (aLPC), inferior frontal sulcus (IFS), posterior medial prefrontal cortex (pmPFC), posterior superior frontal sulcus (pSFS), anterior Insula (aIns), and frontal pole (FP).

Although this task elicits widespread cortical activity, these activated regions include not only those that specifically represent navigation-related information, but also more general areas in the visual and motor cortex that underpin perception and actions. To identify the network of functional regions that specifically represent navigation-related information, we used bootstrapped model connectivity to hierarchically cluster the encoding model weights from all participants on the fsaverage6 cortical surface. This process was used to identify cortical vertices that specifically represent navigation-related information, and to delineate regions of interest (ROIs) on the cortical surface.

This procedure revealed 11 distinct cortical ROIs that represent navigation-related information (Benjamini-Hochberg corrected p < 0.05, permutation test, Fig. 2B, for individual participants see Supplementary figure 5). For ease of reference, in the rest of this paper we refer to these ROIs by their anatomical locations and/or their overlap with ROIs identified in prior studies: PPA-collateral sulcus (PPA-ColS), OPA-posterior lateral parietal cortex (OPA-pLPC), RSC-posterior precuneus (RSC-pPrCu), dorsal precuneus (dPrCu), anterior precuneus (aPrCu), anterior lateral parietal cortex (aLPC), inferior frontal sulcus (IFS), posterior medial prefrontal cortex (pmPFC), posterior superior frontal sulcus (pSFS), anterior Insula (aIns), and frontal pole (FP). The cortical distribution of these ROIs indicates that, during active naturalistic navigation, navigation-specific information is represented in a large functional network that is distributed across the visual, parietal, and prefrontal cortices.

### Functional properties of navigation ROIs

Navigation through the real world is a complex task. Navigational information must be extracted from perceptual inputs, progress must be tracked continuously and compared to internal plans, and motor actions must be produced to move through the world. To quantify the unique functional contributions of each ROI to this complex task, we created a *tuning bias profile* for each ROI that describes how its tuning for all 38 feature spaces differs from that of the rest of the cerebral cortex. Positive values within the tuning bias profile indicate which specific feature spaces are over-represented in that ROI relative to the rest of the cerebral cortex, and negative values indicate which feature spaces are under-represented relative to the rest of the cerebral cortex. Because both human and animal studies have shown that medial temporal lobe (MTL) structures represent cognitive map features ^5,7,43,44^, we also applied this analysis to the hippocampus and entorhinal cortex. Figure 3A-K shows positive tuning bias profiles for each of the 11 navigation-related ROIs identified in this study, along with those for the hippocampus, V1, and M1H (see Supplementary figure 6 for a legend for all colours, and Supplementary figure 7 for negative tuning bias profiles).

**Fig. 3.**
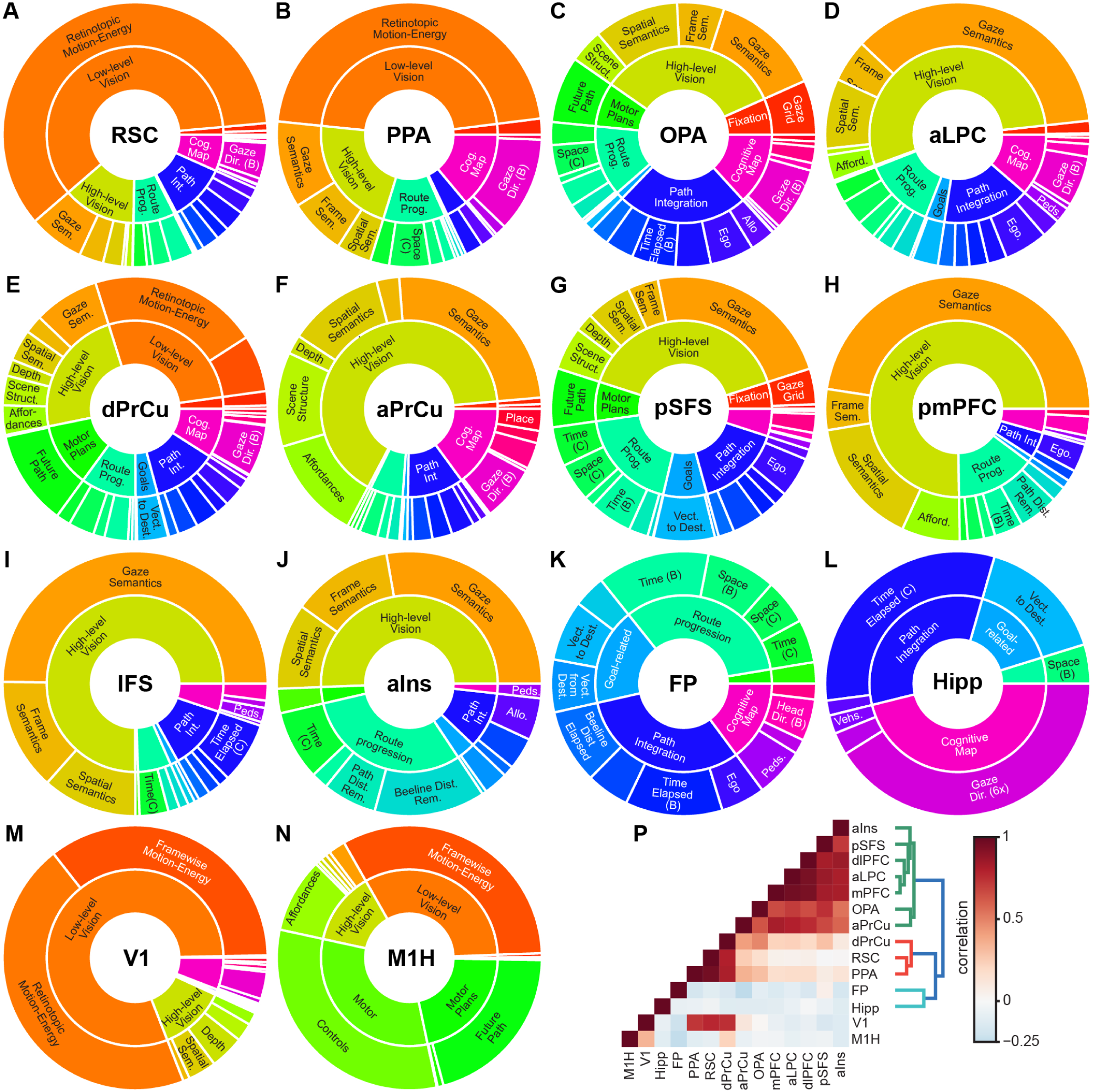
Each component of the navigation network represents a unique combination of navigation-related features. To determine the functional specialization of each of the ROIs within the navigation network, we computed a tuning bias profile for each ROI. The positive tuning biases shown here describe the feature space each ROI over-represents relative to the rest of the cerebral cortex (for negative tuning biases, see Supplementary figure 7). For each ROI, the outer ring in the sunburst plot shows the over-represented feature spaces, while the inner ring shows these feature spaces aggregated together into one of nine categories. These results suggest that each region likely serves as the hub for a specific aspect of spatial navigation. (**A**) The RSC-pPrCu over-represents low-level visual inputs, suggesting that it maps visual inputs to more abstract navigational representations. (**B**) The PPA-ColS over-represents low-level visual inputs, visual semantics, route progression, and cognitive map features, suggesting that it uses visual inputs to identify the scene and objects within it, and also maps these visual inputs to the cognitive map. (**C**) The OPA-pLPC over-represents a mix of visual semantics, motor plans, route progression, path integration, and cognitive map features. This complex combination suggests that this region supports both visuomotor transformations for visually-guided navigation, and also tracks progression along plans and relates it to the cognitive map. (**D**) The aLPC over-represents high-level visual feature spaces, particularly those related to semantics, route progression and path integration feature spaces, suggesting it uses visual inputs to track progression along planned paths and for computing the path travelled. (**E**) The dPrCu over-represents a variety of both low- and high-level visual feature spaces, and also the planned future path of the participant, suggesting that it aids in visuomotor transformations for visually-guided actions. (**F**) The aPrCu over-represents high-level visual feature spaces, particularly those related to the spatial layout of the scene such as depth, spatial semantics, scene structure, and navigational affordances. This combination suggests that the aPrCu represents information useful for immediate navigational actions. (**G**) The pSFS over-represents a combination of high-level visual features, route progression, and path integration feature spaces. This combination and the proximity of this region to the FEF suggests that it produces top-down signals for using visual information to track route progression and for computing the path travelled. The (**H**) pmPFC and (**I**) IFS are heavily biased for visual semantic feature spaces, suggesting that they likely process high-level semantics in the visual scene specifically for navigation-related purposes. J) The aIns over-represents visual semantics and route progression feature spaces, particularly the beeline distance remaining to the goal, suggesting that this region tracks proximity to the goal. (**K**) The FP over-represents route progression, path integration, goal-related, and cognitive map feature spaces, suggesting that it likely represents navigational goals and the plans for achieving them. For comparison, (**L**) and (**M**) show tuning bias profiles for two ROIs that are not part of the navigation network: V1 and M1H. V1 over-represents low-level visual features, while M1H over-represents motor outputs, motor plans, and low-level visual features that are highly correlated with motor actions that affect the optic flow in the stimulus. (**N**) Pairwise correlations between the tuning bias profiles for all navigation-related ROIs and also V1 and M1H. The dendrogram suggests that these navigation ROIs form three clusters. The RSC-pPrCu, PPA-ColS, and dPrcu form one cluster. The aIns, pSFS, IFS, aLPC, pmPFC, OPA-pLPC, and aPrCu form a second cluster. The FP forms a third cluster.

To illustrate this analysis and to verify that it works as intended, we computed tuning bias profiles for two known functional regions: V1 and M1H. Area V1 (Fig. 3M) over-represents low-level visual feature spaces. In contrast, M1H (Fig. 3N) over-represents motor outputs, motor plans, and low-level visual feature spaces that are correlated with motor actions that cause optic flow. These biases agree well with the known functions of these areas, suggesting that tuning bias profiles accurately capture the unique functional properties of these ROIs.

In the anterior visual cortex, we find three regions that overlap with known scene-selective regions. The RSC-pPrCu region has been suggested to be primarily responsible for transformations between reference frames ^45^. We find that the RSC-pPrCu predominantly overrepresents retinotopic motion-energy, a low-level visual feature space, along with some high-level visual, route progression, path integration, and cognitive map feature spaces. The route progression feature spaces were directly adapted from rodent neurophysiology studies ^46,47^ that showed RSC neurons represented route-referenced features. Not only does this result support the reference frame transformation interpretation of RSC’s role in spatial navigation, they also directly replicate rodent neurophysiology findings in human neuroimaging. In addition to the navigation features, the overrepresentation of retinotopic motion-energy features in RSC-pPrCu suggests that it may be a key interface for transforming from retinotopic visual inputs to more abstract navigation-related representations.

The PPA-ColS region has been suggested to be primarily responsible for recognizing the identity of scenes ^19^. We find that the PPA-ColS over-represents low-level visual feature spaces, visual semantics, route progression, and cognitive map feature spaces. The bias for retinotopic motion-energy and visual semantic features agree well with the scene identification interpretation of PPA’s role in spatial navigation. The bias for gaze direction also supports this interpretation, as PPA has been shown to be selective for specific viewpoints within a scene ^18^. However, the bias for route progression representation suggests that during active navigation, the PPA-ColS may also be recruited to integrate bottom-up scene identity information with top-down navigational plans to track progress along routes.

The OPA-pLPC region has been implicated to be primarily responsible for visually-guided movement ^10,19^. We find that the OPA-pLPC over-represents high-level visual features, future path, route progression, path integration, and cognitive map feature spaces. The high-level visual bias includes both visual semantics, scene structure, and spatial semantics, suggesting that the OPA-pLPC contains information about the 3D geometry of the environment. Notably, it is the only anterior visual region whose bias profile contains the future path of the subject, which is referenced in egocentric space. This bias agrees well with the visually-guided movement interpretation of OPA’s role. However, the complex bias profile also includes route progression, path integration, and cognitive maps feature spaces, suggesting that during active navigation, the OPA may also integrate concrete navigational actions with top-down navigational plans to track progress and perform path integration.

The aLPC region overlaps with the superior parietal lobule (SPL). The SPL has been shown to be active when subjects identified exits in mazes ^48^, and more recent studies suggest its involvement in visually-guided navigation ^19,49^. We find that aLPC over-represents high-level visual feature spaces, including visual semantics, spatial semantics, and navigational affordances, along with route progression, path integration, and cognitive map feature spaces. These biases suggest that the aLPC, along with OPA-pLPC, may aid in the production of visually-guided navigational actions and also integrate these actions with top-down navigational plans.

Immediately anterior to RSC-pPrCu, we identify two regions in the precuneus that represent navigation-related information. The precuneus is known to be recruited for spatial updating during self-motion ^50^ and visuo-spatial mental imagery ^51–53^, is associated self-reported action planning ^34^, and may be implicated in the perception of self-position within the environment ^54^. We find that the dorsal precuneus (dPrCu) region (Fig. 3E) predominantly over-represents a mix of low-level and high-level visual features, the future path of the participant, route progression features, path integration features. The high-level visual biases include spatial semantics, depth, scene structure, and navigational affordances, which all represent some aspect of the 3D structure of the environment. These representations agree well with the needs of self-motion-related spatial updating: visual inputs are likely used to perform path integration functions and update one’s position along the planned route. The representation of the planned future path suggests that the dPrCu not only processes spatial updating as one moves, but also prospectively plans self-motion through the perceived scene.

The anterior precuneus (aPrCu) region predominantly over-represents high-level visual features, particularly those related to the 3D structure of the scene, namely spatial semantics, the scene structure, and navigational affordances. These biases suggest that the aPrCu may serve to represent information necessary for planning concrete navigational actions. These representations also agree with the possible role of the aPrCu in self-perception: these structural representations of the world likely anchor the perception of one’s own location and relation to the environment. These results lend support to reports that the precuneus is recruited for spatial updating ^50^, and provides concrete evidence that it represents action planning during navigation ^34^.

In the prefrontal cortex, we report five distinct regions: pmPFC, IFS, pSFS, aIns, and FP. The pmPFC had been reported to be active during self-reported action planning and the expectation of landmarks ^34^ and also at decision points ^21^. We find that the pmPFC predominantly over-represents gaze semantics, spatial semantics, navigational affordances, and route progression features. The dominance of semantics, both in visual space and 3D space, is largely consistent with the expectation and searching for landmarks. The representation of navigational affordances is also consistent with action planning, as navigational affordances constrains the actions the participant can take. While ^34^ reported that self-reported action planning engages both preuneus and pmPFC, here we show a putative difference between the functional roles for the precuneus regions and the pmPFC for action planning: the pmPFC likely represents constraints for possible actions, while dPrCU likely represents the concrete planned actions.

The IFS is found in dorsolateral prefrontal cortex, which previous studies have suggested to be recruited during detours ^21,55^ and spontaneous rerouting ^34^. We find that the IFS predominantly over-represents visual semantics, along with some path integration features. This bias profile appears to be largely orthogonal to detours or rerouting. However, our experiment did not include any environmental factors that required detouring or rerouting. Thus, this difference with the prior literature may be due to differences in task demands. Outside the context of spatial navigation, however, the IFS is known for directing object-based attention ^56^ and for planning ^40^. Thus, the IFS may be responsible for navigation-specific attentional control, and for integrating attended objects with navigational plans. Future experiments that induce more rerouting demands during active navigation, and the development of better navigational attention-related feature spaces may better characterize the function of IFS for active navigation.

The pSFS partially overlaps with the anterior portion of the frontal eye fields, which is known for top-down control of visual attention ^57^. We find that the pSFS over-represents a mix of high-level visual features, the future path, route progression, goal-related, and path integration feature spaces. This region has not been well-characterized in the context of spatial navigation, and it is likely that the pSFS, along with the IFS, directs visual spatial attention during active navigation; however, the pSFS may also serve as a hub for integrating aspects of spatial navigation.

The insula has not been well-studied in the context of spatial navigation, though ^34^ had reported that it is active during self-reported expectation of landmarks, visual inspection of the world, and traffic monitoring. We found that the anterior insula (aIns) over-represents visual semantic features and route progression feature spaces, particularly the time elapsed, path distance remaining, and beeline distance remaining along the path. The biases for visual semantics feature spaces agree with the reported activations of the insula when participants expect landmarks and inspect the world. The additional biases for route progression feature spaces suggest that the insula may be a key overlooked region for active navigation that may use visual inputs to track progression along planned routes, and is a promising target for future experiments.

The frontal pole (FP) had been shown to be selective for the distance to the goal ^58^ and during more difficult navigational choices ^59^. We find that the FP over-represents purely abstract navigation-related features, including route progression, goal-related, path integration, and cognitive map features. This combination suggests that the FP may be a convergence region for many aspects of spatial navigation, whose functional properties could not be fully described by more constrained experiments. Given its role in executive control in managing competing goals ^60^, the FP may flexibly combine and prioritize different information for effective spatial navigation. Future experiments that require more task-switching between different aspects of spatial navigation may better characterize FP’s role in managing the many competing demands of active navigation.

The MTL is known for representing a variety of abstract navigation information, including one’s own location ^5^, goal distance ^21^, homing vectors ^61^, among others. We find that the hippocampus over-represents abstract information, particularly the 6-fold representation of gaze direction, the time elapsed since the start of each trial, and a vector to the destination (Fig. 3L). These results agree well with the known roles of the hippocampus in navigation in the representation of abstract cognitive map features and for route planning. (Note that the tuning bias profile for EC could not be determined in this experiment because of magnetic susceptibility artifacts; see discussion and Supplementary figure 8).

Hierarchical clustering of the tuning bias profiles from the 11 cortical ROIs and the hippocampus (Fig. 3O) reveals that they form three clusters: the RSC-pPrCu, PPA-ColS, and dPrcu form one cluster that over-represents low-level visual features, the aIns, pSFS, IFS, aLPC, pmPFC, OPA-pLPC, and aPrCu form a second cluster that over-represents high-level visual features, navigation features, and action outputs, and the FP and hippocampus forms a third cluster that over-represents abstract navigational information. These results suggest that the navigation network of the human brain may be organized around three aspects of spatial navigation: processing visual inputs, forming abstract navigational goals and plans, and integrating these plans with perception to produce concrete navigational actions.

### Gradient-based organization of cortex

Because active navigation through the real world requires the close integration of sensory, cognitive, and motor processes, the navigation network in the cerebral cortex must interact with the visual and motor networks. To understand the relationship between the navigation network and other functional networks, we used UMAP ^62^ to recover a nonlinear 2-dimensional functional embedding space for all cortical vertices. In this embedding space, cortical vertices form a continuous distribution (Fig. 4A), which maps to continuous gradients on the cortical surface (Fig. 4B). Early retinotopic visual areas, such as V1, V2, and V3, occupy one corner of the distribution (upper left of Fig. 4A). Primary motor ROIs for the hands, feet, and mouth occupy the opposite corner (lower right of Fig. 4A). The 11 navigation ROIs lie midway between the early visual regions and the primary motor regions. An unexpected aspect of this organization is that the FEF (which mediates saccade targeting) and hMT (which is selective for global optic flow) lie between the navigation network and the primary motor regions. This organization likely reflects the fact that drivers saccade towards an intended turn 1-2 s before entering the turn ^63^, and they slow down before turning. Thus, both saccades and changes in optic flow predict upcoming motor actions. In sum, the cortical embedding space reveals that the navigation network acts as the middle component of a perception-decision-action axis: it applies navigation-related cognitive functions to visual representations to make decisions, and then routes action commands to motor representations.

**Fig. 4.**
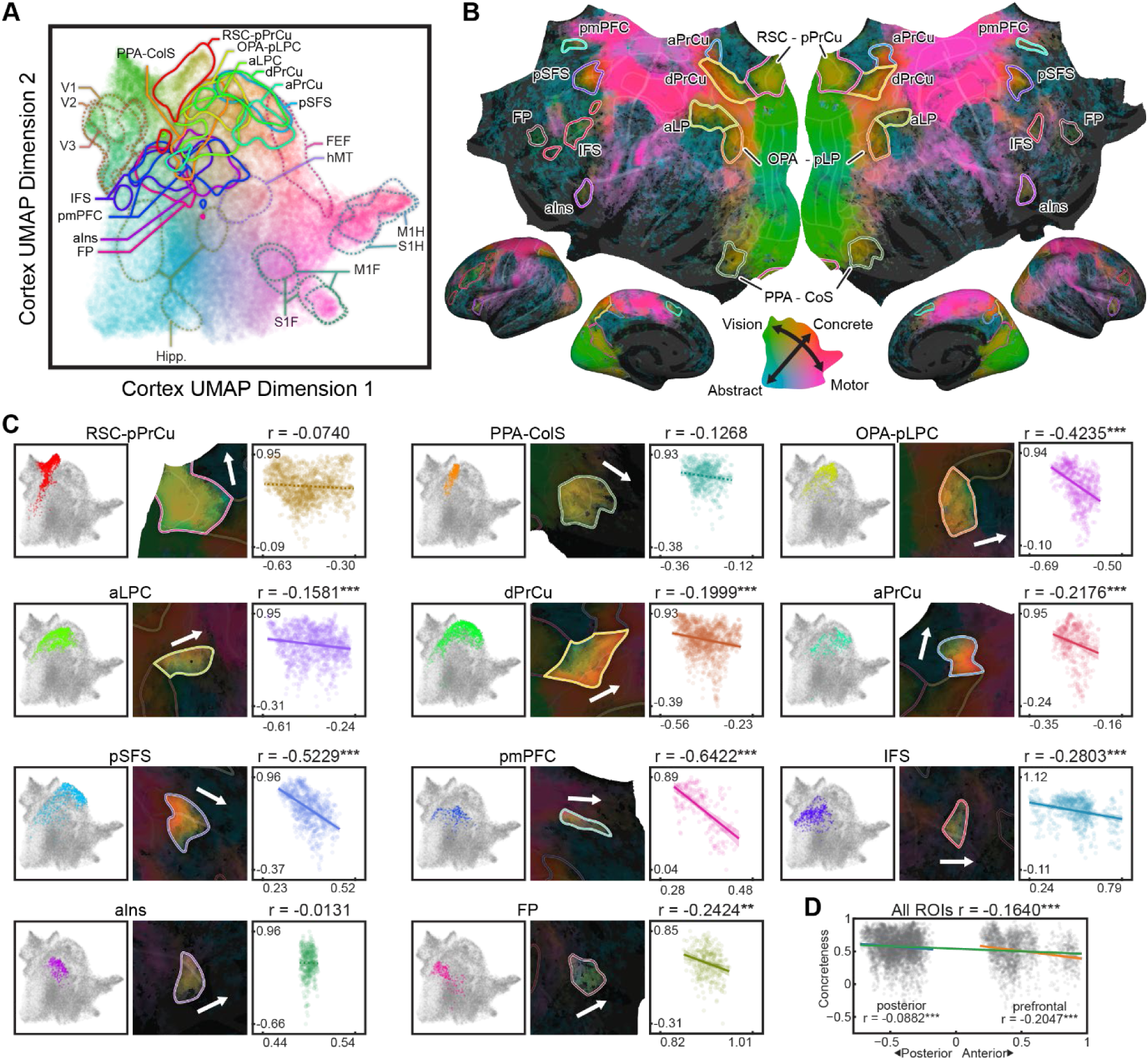
The navigation network is organized along broad functional gradients in the cortex. The navigation network must interact with other cortical systems as we move through the world. A) To understand the functional relationship between the navigation network and other cortical regions, we used UMAP to recover a 2D whole-cortex functional space from the high-dimensional model weights of all vertices. Each vertex was projected to a point in this functional space, and coloured by their location in this space. The locations of the 11 ROIs in the navigation network (solid lines), along with several ROIs from the visual and motor systems (dashed lines), are outlined (contour lines are drawn at the 2.5% density level for each ROI). The 11 navigation ROIs lie between the early visual regions and the primary motor regions, suggesting that the navigation network applies navigation-related cognitive functions to visual representations to make decisions, and then routes action commands to motor representations. B) This functional distribution maps to continuous gradients on the cortical surface. Vertices on the cortex are coloured by their location in the functional space as in (A). C) Detailed views show that functional tuning in each navigation ROI spans a substantial portion of the cortical gradients. Left panels show the location of vertices from each ROI within the functional space, and right panels show the gradient within each ROI. Tuning within each ROI follows broader cortical gradients that span multiple ROIs. Furthermore, in the functional space, the 11 ROIs appear to be organized along a second axis orthogonal to the visual-navigation-motor axis. This axis largely corresponds to a posterior-to-anterior anatomical organization of the 11 ROIs, and suggests this axis may be a gradient between anterior regions that represent more abstract information and more posterior regions that represent more concrete information. These patterns suggest that the navigation network consists of a series of partly redundant functional representations that are organized along broad gradients.

The identification of distinct functional ROIs comprising the cortical navigation network does not necessarily imply that functional tuning within each ROI is completely homogeneous. In fact, inspection of each individual ROI (Fig. 4C, left and middle panels) suggests the contrary: tuning within each ROI appears to be aligned with broader cortical functional gradients that span multiple ROIs (Fig. 4B). Further evidence for this global organization influence can be found by inspection of the cortical embedding space (Fig. 4A). Within this space, the 11 ROIs comprising the navigation network are organized along a second axis that is orthogonal to the visual-navigation-motor axis (Fig. 4A). This second axis coarsely aligns with the posterior-to-anterior anatomical organization of the 11 ROIs. This axis appears to reflect a gradient from more posterior ROIs that represent relatively more concrete information, to more anterior ROIs that represent relatively more abstract information.

To quantify this concreteness gradient, we defined a concreteness index for each vertex based on its split R^2^ scores, and then examined how this index varies posterior-anteriorly across the navigation ROIs. We find that concreteness is significantly correlated with the posterior-anterior position of the vertices across the navigation network. Intriguingly, further inspection indicates that this gradient is replicated within ROIs. Of the 11 ROIs, nine show a significant posterior-anterior concreteness gradient (Fig 4C, right panels). These results not only lend support to the network gradient-based organization of the cortex, but also suggest that these gradients are replicated across multiple regions in a distributed, partially redundant manner.

### Banded ridge effectively disentangles stimulus correlations

The close coordination between perception, cognition, and action during active navigation inevitably induces correlations among the features used to fit encoding models. One concern is that perhaps these stimulus correlations biased our results. Banded ridge regression was specifically developed to account for these correlations during model fitting ^36,37^. To verify that banded ridge was indeed able to account for these correlations, we examined the effects of stimulus correlations on the fit models. Results show that the stimulus correlations do not explain any variance in the model correlations on the cortex (R^2^ = −55.1±14.1, mean ± std.ev. across participants, Supplementary figure 9), indicating that the fit models are unlikely to be driven by stimulus correlations. Thus, these results demonstrate that banded ridge regression effectively disentangles correlated feature spaces during model fitting, and that the results reported here were not unduly biased by stimulus correlations.

### Weak correlations between encoding model performance and behaviour

Because the routes travelled by participants are self-directed, behavioural variability should be reflected in brain activity. To explore whether there are systematic relationships between encoding model performance and behavioural performance, we calculated model prediction performance in individual trials, and summed the split R2 scores across navigation feature spaces. We then tested whether the performance of these navigation models was correlated with path optimality on each trial. This analysis was performed both across the entire cerebral cortex, and also separately within each ROI in the cortical navigation network. To account for differences in SNR and performance across subjects, both the split R^2^ scores and path optimality values were z-scored before aggregation. Because models generally perform better on training data than on test data, we examined this relationship in the train and test sets separately to avoid Simpson’s paradox. No test passed an FDR-corrected p < 0.05 significance threshold. (This likely occurred because subjects’ behaviour was close to the performance ceiling, so there was likely not enough variation in behavioural performance to establish significance.) However, the data show weak, non-significant evidence suggesting that model performance decreases as paths become less optimal in the test set (Supplementary figure 10). These results suggest that perhaps poor behaviour may be reflected in poorer representation of navigation-related information in the brain, but further study will be required to better characterize this relationship.

## Discussion

To fully map the functional network in the human cerebral cortex that mediates spatial navigation, we fit high-dimensional models to brain activity recorded from participants driving in an immersive virtual world. We identified 11 cortical regions across the visual, parietal, and prefrontal cortices that mediate active navigation, each of which represents a unique combination of navigation-related information. Analysis of the model weights show that functional tuning in this network is organized along multiple cortical gradients, and is consistent with results from wide-field neurophysiology studies in rodents ^24,25^, suggesting that the organizational principles of the cortical navigation network may be conserved across species.

These results provide compelling evidence for a distributed network-based, rather than modular, organization of the navigation system in the human cerebral cortex ^3^. Each functional region integrates multiple types of information from multiple levels of processing, and each type of information is conversely distributed across many regions. Each region likely serves as the hub for a specific aspect of spatial navigation. This distributed organization likely reflect feedback and lateral processes in the navigation system at all levels, in addition to feed-forward processing. By identifying the components of the navigational system and their functional properties, these results provide crucial constraints for the development of biologically plausible computational models of human navigation that explain the entire perception-decision-action loop.

Four of the 11 navigation ROIs identified in this study, namely the aLPC, IFS, aIns, and pmPFC, have been identified in previous studies as part of the multiple demand (MD) network ^64^. This overlap between the navigation network and the MD network are not mutually incompatible assignments. The MD network is associated with demanding cognitive tasks. Active navigation in a naturalistic world is a challenging task that requires close integration of perception, action, and multiple cognitive processes, and requires planning across multiple timescales. Therefore, it is unsurprising that the cortical navigation network may recruit MD regions. While these regions are a part of the cortical navigation network, this labelling does not imply that navigation is the sole function of these regions.

One prior experiment also used an active taxi driver task in a naturalistic virtual world ^34,65^. However, that study only produced maps for various subjective self-reported mental states during driving. While that earlier study could show that different mental states corresponded to activation in different brain regions, it could not describe what information is represented within each brain region, or how those regions are functionally organized to support active, naturalistic navigation. Here, we used state-of-the-art high-dimensional encoding model methods to create data-driven, quantitative maps that reflect precisely how navigation-related information is represented across the cerebral cortex.

The tuning bias profile analysis used here failed to identify any significant tuning biases in the EC. However, this should not be taken as a definitive result. The EC is strongly affected by magnetic susceptibility artifacts that reduce signal-to-noise ratio (SNR) ^66,67^. Some prior studies targeting the EC have used EC-optimized MR sequences ^68^, but such MR sequences reduce signal elsewhere in the cerebral cortex. Because we aimed to recover the navigation network across the entire cerebral cortex, we were unable to use this strategy, and so the SNR in MTL structures and especially in EC was quite low relative to other parts of the cortex (see Supplementary figure 8). Future experiments optimized to recover signals in the EC may offer more insights on the functional properties of the human EC during naturalistic navigation.

One limitation of our study is that the supine position of the participants in the MRI scanner eliminates vestibular signals that normally occur during real-world navigation. This limits the ecological validity of the experiment somewhat ^69^. However, behavioural evidence shows that humans preferentially use visual cues over proprioception for navigation ^70,71^. Further, our route progression models replicate neurophysiology findings from freely running rodents ^46,47^; this replication across imaging modalities, species, and experiments suggest that the lack of vestibular signals did not unduly bias our results.

The accompanying online viewer will allow for in-depth exploration of the functional maps produced here. These functional maps provide rich targets for future experiments on active navigation, such as more in-depth studies on the functional properties of each of the cortical navigation regions, or the construction of end-to-end computational models of human spatial navigation. These maps also provide a detailed baseline for studying how the navigation network changes with learning, and may provide a sensitive biomarker for quantifying cognitive changes due to neurodegenerative diseases that affect navigation. Finally, the distributed and partially redundant organization of the human cortical navigation network may provide guidelines for architecting autonomous vehicles that are robust, flexible, and independent.

## Methods

### Participants

Six healthy adult volunteers (3 female, ages 24-34) with normal or corrected to normal vision participated in this study. Participants were recruited from the UC Berkeley student and staff population, and included two of the authors (T.Z. and E.M). All participants had some experience with video games, though the amount of experience, the type of video game, and familiarity with controls varied greatly. All participants extensively practiced with the driving simulator and were familiar with this particular software before scanning.

The experimental procedures were approved by the Institutional Review Board at the University of California, Berkeley, and written informed consent was obtained from all participants.

### Driving simulator for active navigation

We used Unreal Engine 4 (Epic Games) and the Carla plugin ^32^ to build a 2×3 km virtual city (https://github.com/gallantlab/Driving-Simulator). This city contains multiple neighbourhoods, including urban, commercial, suburban, and rural areas, each with a distinct visual appearance. The map contains over 200 uniquely identifiable locations. This city is populated by AI vehicular and pedestrian traffic. AI vehicles obey traffic lights and speed limits, and respect pedestrian right-of-way. This provides an ecologically valid, dynamic world in which participants can actively navigate by driving a virtual car. Video is projected onto a screen in the bore. Participants are shown a first-person view from the virtual car, and drive using a set of MR-compatible steering wheel and pedals. Participants can also use a button to toggle between forward and reverse gears.

The driving simulator produced a complete record of the experiments. This record included ground truth about the local environment from the participant’s point of view, the control inputs and behaviour of the participants, the navigational goal of the participant, and the behaviour of all other vehicles and pedestrians. Additionally, eye motion, respiration, and heartbeat were recorded continuously to facilitate data analysis and modelling. These records were then used to extract features from the experiments offline.

### Task

Prior to scanning, the simulator software was distributed to participants, and they used it to learn the layout of the map and the locations of all possible destinations. During this learning phase, the simulator directed the participants to visit every location and explore the map. To assist in learning, the simulator displayed on-screen navigation information including street names, compass heading, directions, distance to the active destination, and a top-down minimap.

During each MRI session, participants performed a taxi-driver task ^34^ in the virtual world. Of the 200-plus locations in the world, 77 were selected as destinations for the experiment. These locations were distributed across the map and included landmarks (e.g. a statue of a fire hydrant), special named locations (e.g. the Midtown Bank), house numbers (e.g. 1 Broadway), and street intersections (e.g. Market St & Atlantic Ave). As in real-world environments, urban areas contained the densest concentration of destinations, followed by the suburban areas and the rural areas.

Each MRI session consisted of many trials in which the participants navigated to a destination. At the beginning of each trial, a destination that was at least 120 m away from the participant was randomly selected (mean ± std.ev. distance = 474 ± 320 m, max distance = 1,887 m). The same random seed was used across participants to select destinations, but due to the emergent chaos from this task, subjects did not visit the sequence of destinations. A text cue was displayed at the centre of the screen for 2 seconds. Participants then drove to the destination via the quickest path while obeying all traffic laws. At the destination, Participants were instructed to stop completely to indicate that they had arrived (merely passing by the destination did not constitute arrival). A text cue was then displayed for 2 seconds to acknowledge their arrival. After a randomized jitter of 4-12 seconds, a new trial began. Participants navigated at their own pace and responded to the traffic signals, vehicular traffic, and pedestrians. In pilot experiments, we found that it was difficult to drive while fixating; thus, participants were allowed to move their eyes freely during the experiment.

### Scanning procedure

MRI data were acquired on a 3T Siemens Trio with a 32-channel head coil, located at the University of California, Berkeley. BOLD data were acquired using a T2*-weighted gradient-echo EPI sequence customized with a water-excitation radiofrequency pulse to prevent contamination from fat signal (TR = 2.0045 s, echo time = 34 ms, flip angle = 74°, voxel size = 2.24 × 2.24 × 3.5 mm^3^, field of view = 224 × 224 mm^2^, matrix size = 100 × 100, and 30 axial slices to cover the entire cortex). Custom personalized headcases were used to stabilize the head and to reduce motion artifacts ^72^. Before each functional run, a GRE fieldmap was collected for distortion correction. To facilitate cortical surface reconstruction, anatomical data were also collected (three-dimensional T1-weighted MP-RAGE sequence, 1 × 1 × 1 mm^3^ voxel size and 256 × 212 × 256 mm^3^ field of view).

Data were collected across multiple scanning sessions. Each session consisted of six 11-minute runs. Pilot experiments showed that good voxelwise encoding models could be fit with about two hours of data, so at least two hours of data were recorded from each participant (2 hours in P1 and 3 hours each from P2-6).

Respiration and heart rate were recorded using a BIOPAC MP150 system (BIOPAC Systems, Inc.). Eyetracking data were collected using an Avotec dark-pupil IR eyetracker at 60 Hz. The video data were processed with custom software to extract gaze locations (https://github.com/gallantlab/Eyetracking). To ensure accurate eyetracking calibration, at the beginning of every 11-minute functional run, 35 calibration points were presented for 2 seconds each (accuracy: 1.2 ± 0.3 degrees of visual angle, mean ± std. across participants). The taxi-driver task began immediately after the eye calibration sequence was completed.

### Behavioural Analysis

A path optimality value was used to quantify the behavioural wayfinding performance of participants. To calculate the path optimality on each trial, A* search ^35^ was first used to find an optimal path between the starting position of the participant and the destination. The path optimality was then defined as the length of the actual path taken by the participant and the length of the A* path. A path optimality of 1 indicates that the participant was optimal in wayfinding; a path optimality greater than 1 indicates that the path taken by the participant is suboptimal. Because of the heuristics that guided A* search in our environment, the A* path length is not guaranteed to be globally optimal; in these rare cases, it is possible for the path optimality to fall below 1.

### fMRI data preprocessing

Each functional run was first motion-corrected using the FMRIB Linear Image Registration Tool (FLIRT) from FSL 5.0 ^73,74^. Next, functional images were unwarped by applying FUGUE from FSL to fieldmaps collected between functional runs. All volumes in the run were then averaged across time to obtain a high-quality template volume. To align data collected across multiple sessions and runs, the template volume from the first session was selected as a target, and the template volume from all other runs across all sessions were aligned to this target. Pycortex ^75^ was then used to align the functional runs to the anatomical surface. Alignment was checked manually and adjusted as necessary to improve accuracy. Low-frequency voxel response drift was identified using COMPCOR ^76^ and removed from the signal. Physiological signals from respiration and heartbeats were regressed out with RETROICOR ^77^. Voxel activity in each 11-minute run was z-scored separately; that is, within each run, the mean response for each voxel was subtracted and the remaining response was scaled to have unit variance. To remove confounds from the eyetracking calibration sequence and detrending artifacts, the first 35 and last 5 TRs were then discarded from each run.

### Cortical surface reconstruction

Freesurfer^78^ was used to reconstruct cortical surface meshes from the T1-weighted anatomical volumes. The freesurfer anatomical segmentation was checked by hand, and Blender (Blender Foundation) and pycortex were used to manually correct the segmentation where necessary. Blender and pycortex were then used to remove the medial wall, and relaxation cuts were then made into each surface. The cut at the calcarine sulcus was made using retinotopic localizers as a guide to bisect V1 along the horizontal meridian.

### Voxelwise modelling with banded ridge regression

We used banded ridge regression to fit voxelwise encoding models to the BOLD activity recorded in this experiment. The fit encoding models were used to recover functional maps reflecting how navigation-related features are represented across the cerebral cortex. The voxelwise encoding modelling (VEM) framework is a powerful approach that has been validated in many previous experiments ^79–82^. Conceptually, the time series of the activity in each voxel is modelled as a combination of the timeseries of stimulus and task features, and model weights describe how each feature affects the activity in each voxel. Here, we fit linear encoding models of the form

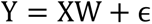

in which Y is the measured brain activity, X are stimulus and task features, and W are weights that describe the relation between features and brain activity. The weights W are estimated by linear regression. In typical cases, ridge regression is used to account for correlations between features. Here, to account for the substantial difference in the number of features across different feature spaces, banded ridge regression ^36,37^ is used to regularize feature space independently. The optimal weights are found by solving the optimization problem

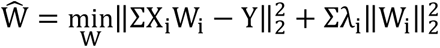

In which X_i_ denotes separate feature spaces, and the optimal combination of λ_i_ are found through cross-validation. We note that mathematically, these voxelwise encoding models are a special form of the general linear model, a statistical framework that is also used by the more traditional statistical parametric mapping (SPM)^83^ framework in neuroimaging, and in the supplementary methods we provide a more detailed discussion comparing the VEM and SPM frameworks.

### Features

In the VEM framework, features are first extracted from the stimulus and task prior to model fitting. Each set of features, or feature space, is a nonlinear embedding of the stimulus or task that explicitly extracts some information. Each feature space can be conceptualized as a hypothesis of what information the brain might represent. In this experiment, both the animal and human navigation literature were used to define many different navigation-related feature spaces. Several other feature spaces were also included that we hypothesized might be navigationally informative, but which have not been investigated previously. In total, we constructed 38 feature spaces that total 28,134 features. (A brief description of each feature space can be found in the supplementary text.) All feature time series were first computed on a frame-by-frame basis, and then downsampled to the fMRI sampling rate ^84^. Together, these feature spaces encompass many aspects of navigation, including perceptual information such as the motion-energy content of the stimulus, abstract navigational information such as a vector to the destination, and motor actions such as the steering wheel angle.

### Model fitting

Models for all feature spaces were simultaneously fit to the BOLD activity of each voxel independently. To account for differences in the size of different feature spaces, banded ridge regression ^36,37^ was used to impose a different regularization parameter on each feature space separately. The hemodynamic response function for each voxel was captured by a finite impulse response (FIR) filter with five delays. The optimal feature weights and the shape of the FIR filter for each voxel were empirically estimated by cross-validation. During fitting, 42,000 random combinations of regularization parameters were explored. Model prediction accuracy was quantified by the coefficient of determination (R^2^) between the predicted and actual brain activity on a hold-out dataset not used during model fitting. Permutation tests were used to estimate statistical significance in each voxel. A significance mask was determined at a Benjamini-Hochberg FDR corrected p < 0.05 level. A group-level significance mask was then determined by selecting vertices on the fsaverage surface where at least half the participants had significant predictions. For each voxel, the *joint model* performance of all 38 models was partitioned across the models for each of the 38 feature spaces. We term these values the *split R*^2^ *scores*.

### Effects of correlated features

A major complication in the analysis of data from naturalistic tasks is the inherent correlation between features. Banded ridge regression was developed to address this issue mathematically during model fitting by independently regularizing feature spaces ^36,37^ and searching through the hyperparameter space of regularization parameters for an optimal model fit. Furthermore, banded ridge also accounts for variance explained by correlated feature spaces when computing split R^2^ scores to partition model performance across feature spaces.

To verify that banded ridge regression was indeed able to account for these correlations, we quantified their relationship with the patterns of results we report in our study. For each pair of feature spaces, we calculated both the feature correlations and their model correlations. Feature correlations were calculated as the ordinary least squares regression prediction performance between the two feature spaces. Features from each feature space were used as regressors to predict the time series of the other features, and the mean correlation was taken across all features within each pair of feature spaces. The model correlations were calculated as the correlation between the prediction performances across the cortical surface for the encoding models for the two feature spaces. The variance explained by the feature space correlations in the fit model correlations were computed to determine whether stimulus correlations drove the relationships between the fit models.

### Noise ceilings

In experiments with fixed or more controlled stimuli, it is possible to repeat trials multiple times to estimate noise. However, in an interactive experiment such as driving, it is impossible to repeat trials. Although the simulator can be configured to present the same situations, participants cannot reliably perform the same action sequence identically. Because the virtual world is a complex dynamical system, small deviations will cause conditions to quickly diverge across repeats. Because of this lack of repeats, we cannot estimate a functional noise ceiling in these data. Thus, all model prediction accuracies are presented without noise ceiling correction.

### Model weight normalization

The fit encoding model weights describe how each feature contributes to the activity in each voxel. Thus, the organization of the functional networks can be recovered by finding voxels with similar model weights. However, the banded ridge regression procedure optimizes the regularization hyperparameters separately for each voxel. While this procedure maximizes model performance in each voxel, it produces model weights with different scales across voxels. The scaling difference makes it difficult to compare weights across voxels to recover functional maps.

To remove the scaling differences, the weights for each model in each voxel were first normalized to have unit norm. Next, the prediction accuracy of each model was normalized across all significant voxels such that the 1^st^ percentile became 0 and the 99^th^ percentile became 1. The model weights were then rescaled by this normalized prediction accuracy. This rescaling effectively corrects for the differences in model weight scales induced by different regularization hyperparameters between voxels.

### Identifying the navigation network

To recover the functional cortical network mediating naturalistic navigation from the high-dimensional encoding models, we applied model connectivity ^38^ to the fit models, and bootstrapped the model connectivity solutions.

#### Model connectivity

Briefly, FreeSurfer’s mri_vol2surf was first used to project model weights and significance masks from individual participants to vertices on the fsaverage6 surface (the fsaverage6 surface was selected because its number of vertices best approximate the number of voxels in the data). Vertex-wise model weights were then averaged across participants to obtain a group-level weight map. The group-level significance mask was selected as the vertices where at least half of the participants had significant predictions. While this choice is somewhat arbitrary, it has no practical effect on the results for two reasons. First, the masking occurs in vertices with negligible weights that are ignored by the MC process, and therefore the exact choice of threshold does not impact the MC solution; the masking rather serves to reduce computational complexity for the clustering algorithm by reducing the number of samples. Second, the bootstrapping process (see next section) varies the specific masks used across bootstraps, and therefore the final result is not sensitive to the mask choice. A hierarchical clustering linkage matrix was then computed for all significant vertices based on their weight similarities. This linkage matrix was then used to determine a clustering solution for a desired number of clusters.

During model connectivity estimation, several hyperparameters must be chosen. These hyperparameters include the distance metric used to determine the similarities between vertices and the number of distinct networks into which to group the vertices. Furthermore, because the feature spaces used in this experiment vary greatly in size, the distance estimate between vertices may be biased towards higher-dimensional feature spaces. To account for this bias, for each feature space with a dimensionality higher than 10, we calculated the first 10 principal components of its model weights. Thus, the hyperparameters also include a choice between the full weights and the reduced dimensionality weights. The final model connectivity clustering solution is dependent on all these hyperparameters.

#### Bootstrapping and identifying the cortical navigation network

Because the exact model connectivity clustering solution is dependent on a variety of hyperparameters, we bootstrapped the model connectivity process to obtain a stable estimate of the extent of the cortical navigation network. First, 200 separate hierarchical clustering linkage matrices were computed. On each iteration, a random combination of distance metric (correlation or Euclidean distance) and model weights (full weights or reduced dimensionality weights) were chosen. The combination of subjects and weight dimensions were then bootstrapped. A linkage matrix was then computed for this bootstrapped data. Then, from this pool of 200 bootstrapped linkage matrices, 1000 random clustering solutions were generated. For each solution, a random number of desired clusters between 4 and 16384 was selected, and a random linkage matrix from the pool of 200 linkage matrices was used to calculate a model connectivity clustering solution.

With this set of bootstrapped model connectivity clustering solutions, we defined a navigation preference index (NPI) for each vertex as the frequency that it was grouped into the cluster with the highest average split R^2^ score for models for navigation feature spaces. The cortical navigation network was then defined as the collection of vertices with high NPIs. Permutation tests were used to establish the significance of the NPI at each vertex. On each iteration, the clustering assignments are randomized across the cortical surface to establish a null distribution for the NPI. This process was repeated 1000 times. The navigation network was identified as the set of vertices with high and significant NPIs. Variations in the NPI across the cortical surface were used to define boundaries between adjacent ROIs.

### Tuning bias profiles

The split R^2^ scores quantify how the representation of each of the 38 feature spaces contributes to the activity at each vertex. These split R^2^ scores were averaged across vertices within each of the 11 navigation ROIs to describe their respective functional properties (Supplementary figure 11). To highlight the functional properties of each ROI, we calculate how its split R^2^ scores differed from the rest of the cortex. Thus, the difference in the average split R^2^ score within an ROI against that for the rest of the cortex quantifies the features that the ROI over- and under-represents relative to the rest of the cortex. We term this difference the tuning bias profile for each ROI. Permutation tests were used to establish the significance of the tuning bias profiles. On each iteration, the ROI assignments were randomized, and a null tuning bias profile was computed for each ROI. This process was repeated 1000 times, and we report only the significant tuning biases within each ROI.

### Visualizing the functional organization of the cortex

To visualize the functional organization of the cortex, we applied UMAP ^62^ to the high-dimensional encoding model weights. Because the weights for all 28,134 features are high-dimensional and non-gaussian, linear dimensionality reduction methods such as PCA are ill-suited for visualization (Supplementary figure 12). To reduce the influence of feature space size biasing towards higher-dimensional feature spaces, we used the top 10 PCs of feature spaces with a dimensionality higher than 10. Because UMAP results are dependent on hyperparameters, we explored many combinations of UMAP parameters. In the final projection, we chose 256 neighbours and a minimum distance of 0.5. Because UMAP is also stochastic, we ran repeated instances of UMAP with the same parameters to ensure that the recovered space is stable.

### Concreteness index

Inspection of the functional organization of the cortex suggests that, in addition to a visual input-to-motor output gradient, there is an apparent posterior-anterior gradient of concreteness of the representation. To quantify this gradient, we used the split R^2^ scores to define a *concreteness index* (CI) for each vertex. We operationally defined concrete feature spaces as those that can be determined from the visual scene alone and the actions produced by the participants. These corresponded to the feature spaces in the fixation-related, low-level visual, high-level visual, and motor categories. Abstract feature spaces were defined as all other feature spaces. For each vertex, the CI is defined as the sum of the split R2 scores for the concrete feature spaces, minus the sum of the split R2 scores for abstract feature spaces, normalized by the total R^2^. The CI is on a range of [−1, 1], in which −1 corresponds to a representation of only abstract feature spaces, and +1 corresponds to a representation of only concrete features.

For each vertex, we also calculated its surface-based posterior-anterior position. To do so, we first identified the anterior-most and posterior-most vertices in volumetric space. Then, for every other vertex, we calculated its geodesic distance along the cortical surface to these two vertices. The posterior-anterior position for each vertex was then defined as the distance to the anterior vertex minus the distance to the posterior vertex, divided by their sum. This position is on a range of [−1, 1], in which −1 is the posterior-most vertex, and +1 is the most anterior vertex. The CI and posterior-anterior positions were then correlated to quantify this concreteness gradient.

## Data availability

The datasets collected and analysed during the current study will be made available on GIN (https://gin.g-node.org) upon publication.

## Code availability

The driving simulator and associated code will be made publicly available online at http://github.com/gallantlab/driving-simulator, http://github.com/gallantlab/driving-launcher, and http://github.com/gallantlab/driving-utilities upon publication. Analysis code will be made publicly available at http://github.com/gallantlab/driving-network upon publication.

## Acknowledgments

We thank all members of the Gallant Lab for discussion and support throughout this work, especially A. Nunez-Elizalde, M. Lescroart, J. Gao, M. Eickenberg, S. Slivkoff, and S. Popham. We also thank Dimitar Filev and Ken Washington at the Ford Motor Company, and Marc Steinberg at the Office of Naval Research, who provided critical funding support during the early stages of this project.

## Funding

National Institutes of Health grant R01-EY031455 (JLG)

Office of Naval Research grant N000142012002 (JLG)

Ford University Research Program (JLG)

Office of Naval Research DURIP awards N00014-22-1-2217 (JLG)

Office of Naval Research Multidisciplinary University Research Initiative grant N000141410671(JLG)

National Science Foundation Graduate Research Fellowship Program DGE 1106400 & DGE 1752814 (TZ)

## Author contributions

Conceptualization: TZ, JLG

Methodology: TZ, JLG

Investigation: TZ

Formal Analysis: TZ, EM

Writing – original draft: TZ

Writing – review & editing: TZ, JLG, EM

