## Supplementary figures, table, and methods for "A map of the cortical functional network mediating naturalistic navigation"

**The file includes:**

Supplementary figures 1-12
Supplementary table 1
Supplementary methods

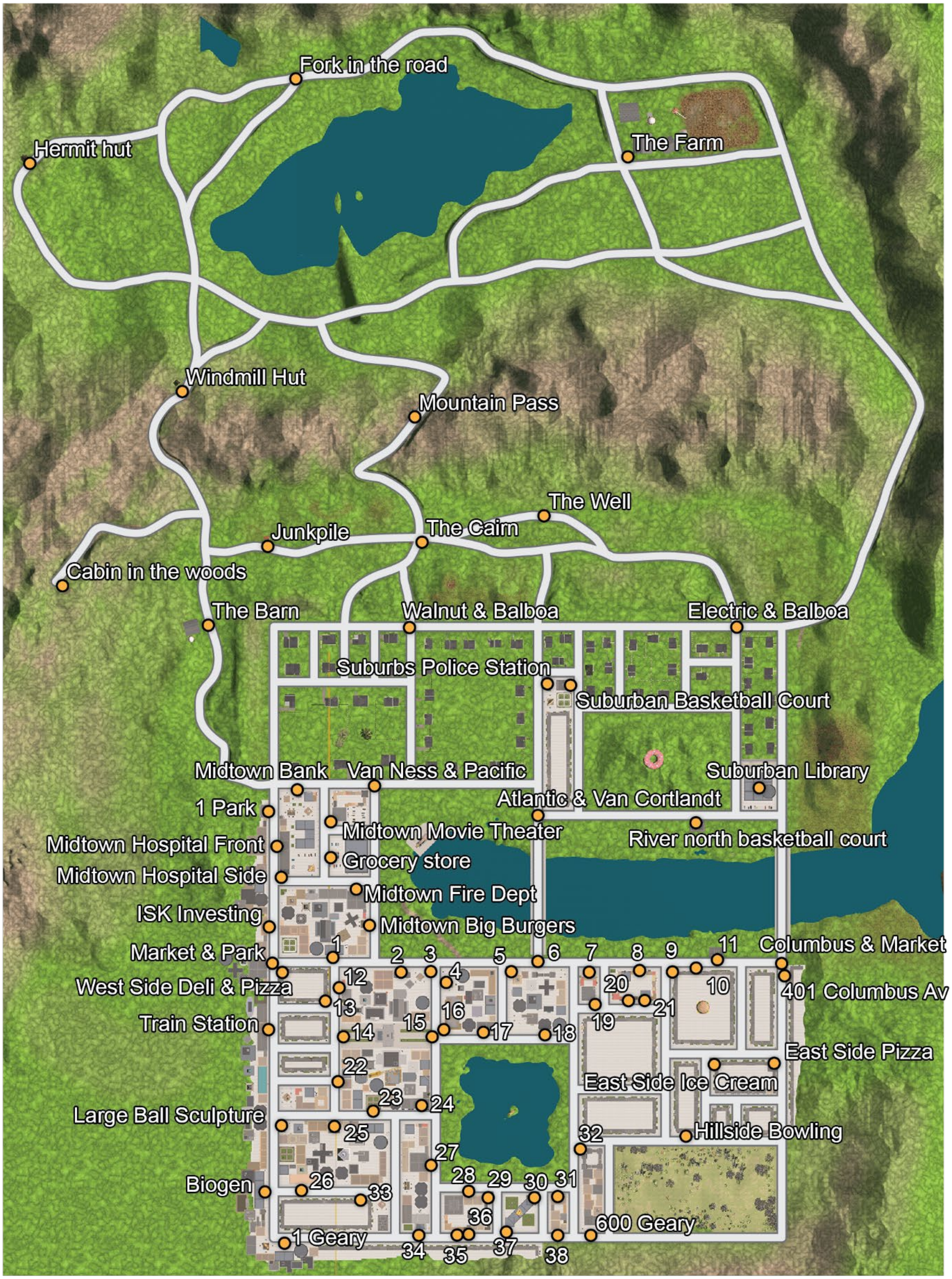


1. Southwest Holdings
2. Downtown Police station
3. Downtown Donuts
4. Pizza On Canal
5. Downtown Bank
6. Market & Atlantic
7. Riverfront Eats
8. Riverfront Cafe
9. South River Corner Cafe
10. South River Noodles
11. South River Basketball Court
12. 1 Broadway
13. Broadway Big Burgers
14. City Hall Front
15. City Hall Back
16. Downtown Construction Site
17. 200 Lake Park N
18. Inicorp
19. South River Movie Theater
20. South River Bar
21. South River Music Store
22. Downtown Library
23. Downtown Minipark
24. Whitehall Hotel
25. Downtown Music store
26. Downtown Hospital
27. Lake Park W Fire Dept
28. 3 Lake Park S
29. Lake Park S Police Station
30. Trousers North Tower
31. Lake Park S Bank
32. Lake Park E Police Station
33. Southtown Noodles
34. Museum Front Entrance
35. Southtown Bank
36. Southtown Pizza
37. Trousers South Tower
38. Park South Library

**Supplementary figure 1. Map of the virtual city with all navigation targets.** Orange dots indicate all possible destinations for the taxi-driver task.

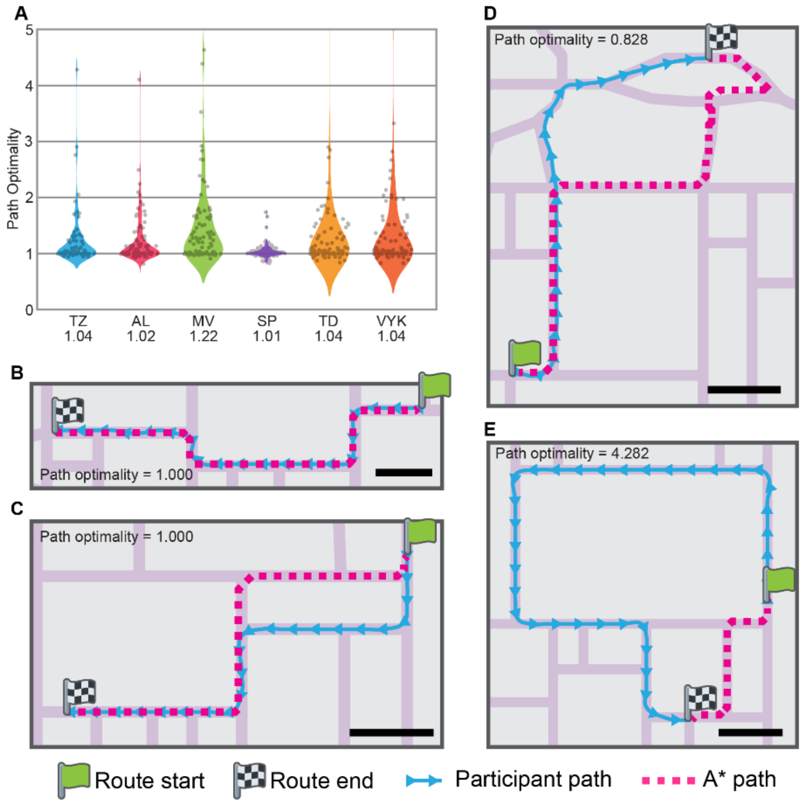


**Supplementary figure 2. Participants were near-optimal in pathfinding.** To quantify the behavioral performance of the participants, we defined a path optimality metric for each trial as the ratio between the length of the actual path taken by the participant and the length of a path found by A* search. A) Path optimality for all participants. Each dot represents the path optimality for a single trial. The median path optimality is shown for each participant below the plot. B-E) Paths from example trials. Green and checkered flags respectively mark the start and end routes, and blue and pink lines respectively mark the path chosen by the participant and A*. Arrows on the blue line indicate the facing direction of the subject. Light purple regions indicate roads, and the black scale bar denotes a length of 100 m. B) A trial with a path optimality of 1, in which the participant chose a path identical to that found by A* search. C) A trial with a path optimality of 1, but in which the participant followed a path different to that found by A*. Because of the grid nature of the map, the participant path had the same length. D) A trial with a path optimality less than 1. Because A* is a heuristic-based algorithm, it occasionally fails to find the globally optimal path. In such rare cases, the globally optimal path followed by the participant results in a path optimality less than 1. E) A trial with a path optimality much larger than 1. Because A* does not account for direction-of-travel along a road or for traffic conditions, some A* paths could not be legally followed. Therefore, in some cases, subjects followed an apparently extremely suboptimal path, resulting in a path optimality much larger than 1. In this example, the A* path begins opposite the heading of the participant; in reality, the participant must continue travelling along their heading and drive around a large city block.

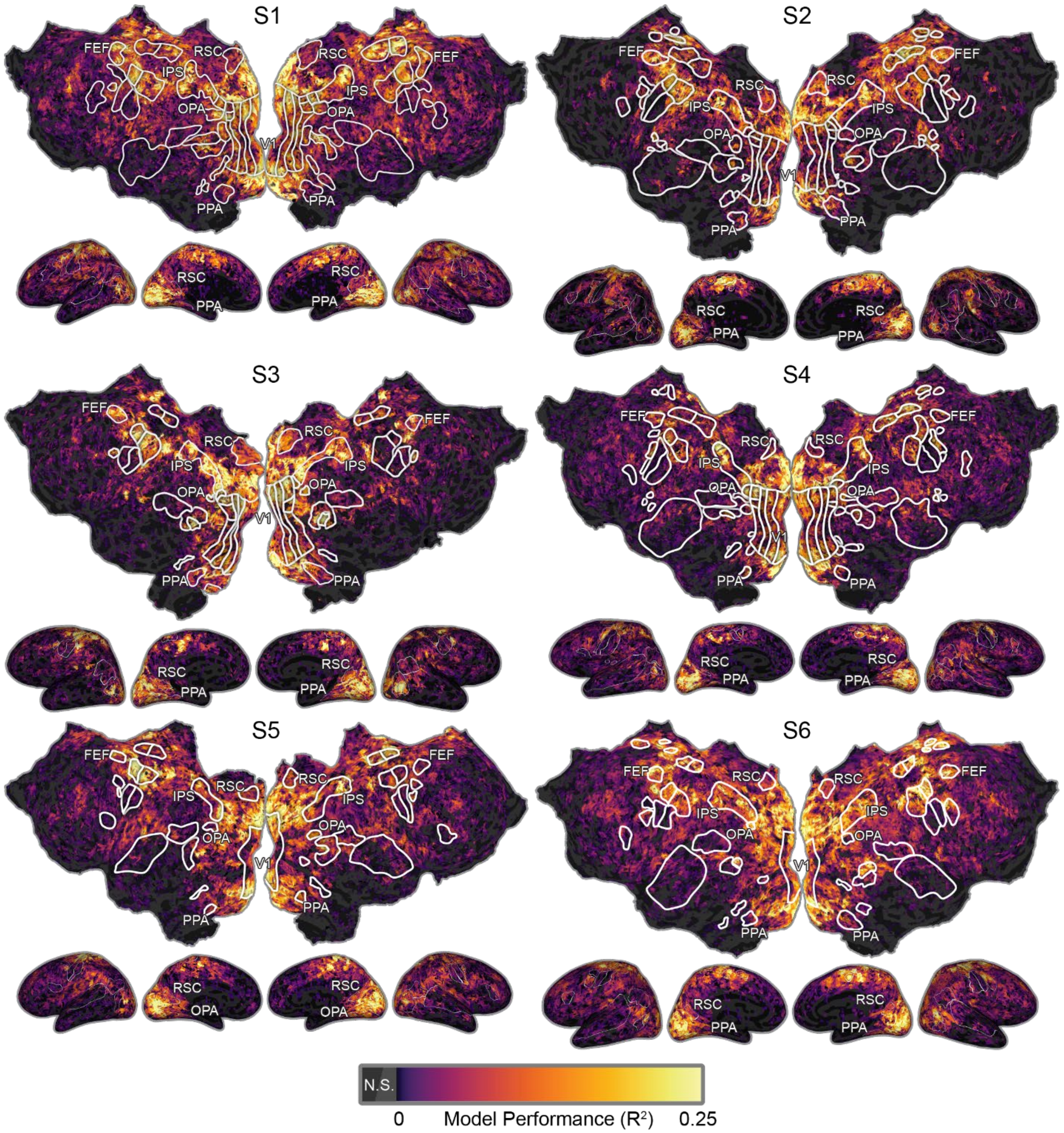


**Supplementary figure 3. Joint model prediction performance in individual participants.** Color scale is the same as Figure 2A. Prediction performance was computed as the fraction of variance explained in each voxel (R2). Voxelwise encoding models predict BOLD responses consistently across participants in many brain areas, including RSC, OPA, PPA, IPS, parietal cortex, Supplemental motor cortex, and DLPFC. Many of these regions have previously been implicated in representing navigationally relevant information in the human brain.

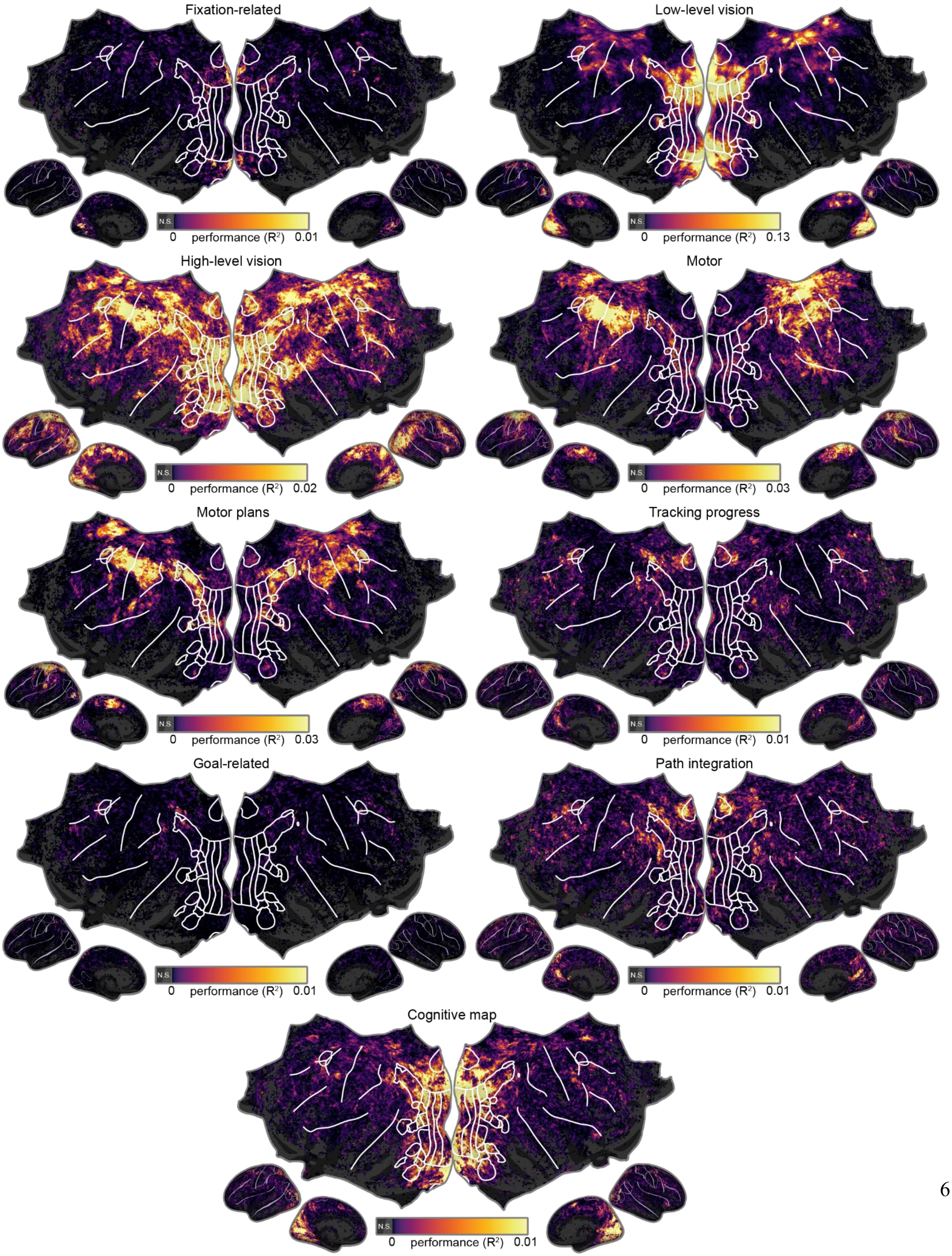
Fixation-related
N.S.
0 performance (R²) 0.01
Low-level vision
N.S.
0 performance (R²) 0.13
High-level vision
N.S.
0 performance (R²) 0.02
Motor
N.S.
0 performance (R²) 0.03
Motor plans
N.S.
0 performance (R²) 0.03
Tracking progress
N.S.
0 performance (R²) 0.01
Goal-related
N.S.
0 performance (R²) 0.01
Path integration
N.S.
0 performance (R²) 0.01
Cognitive map
N.S.
0 performance (R²) 0.01

6

**Supplementary figure 4. Prediction performances for models from each of the nine categories.** The split $R^2$ scores are summed for all models within each category and shown on the cortical surface.

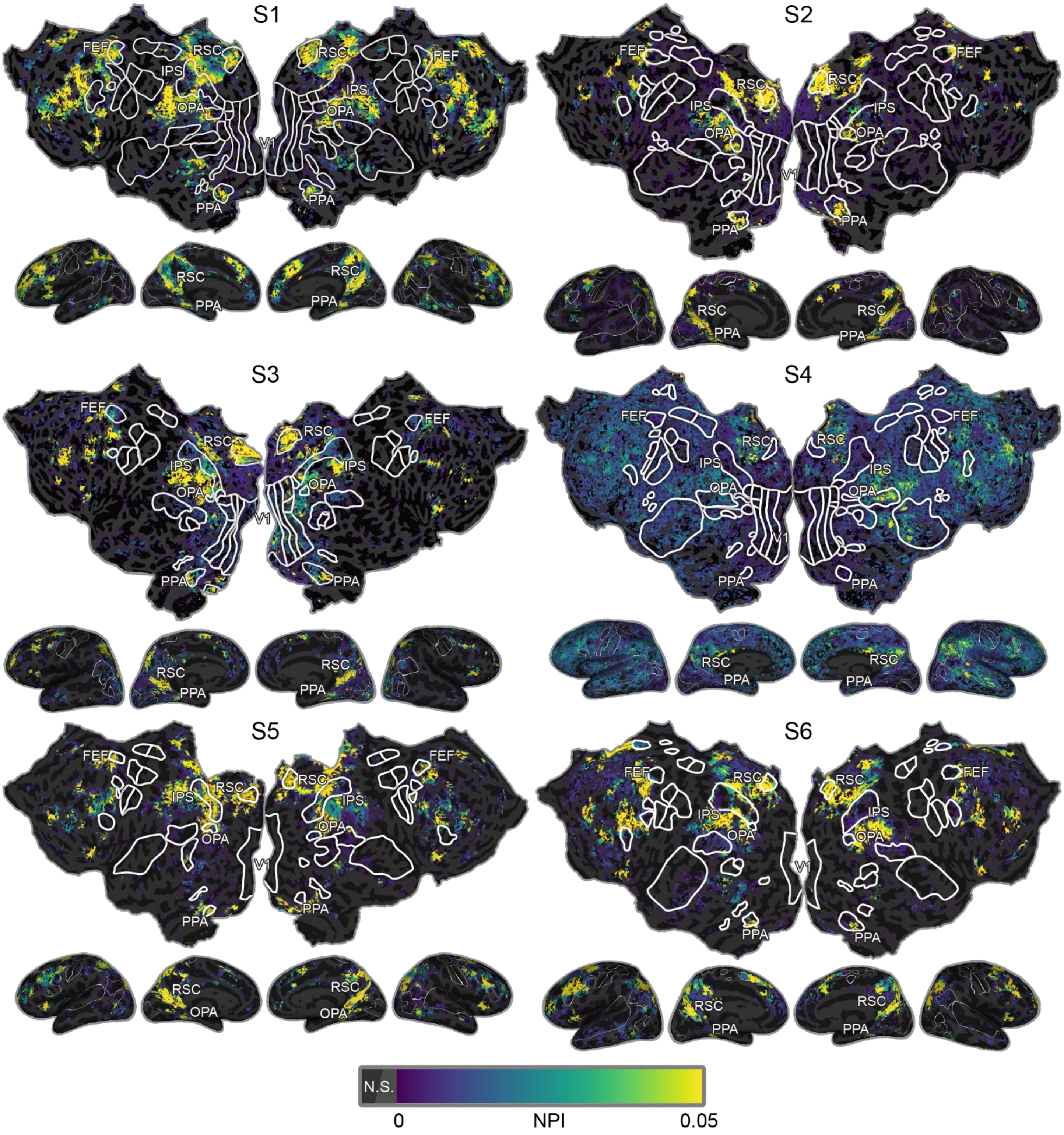


**Supplementary figure 5**. **Navigation preference indices calculated for individual participants.** The same methods for Fig. 2B was used to calculate NPIs within each individual participant. Regions of high NPIs are found across the anterior visual cortex, parietal cortex, and prefrontal cortex in individual subjects.

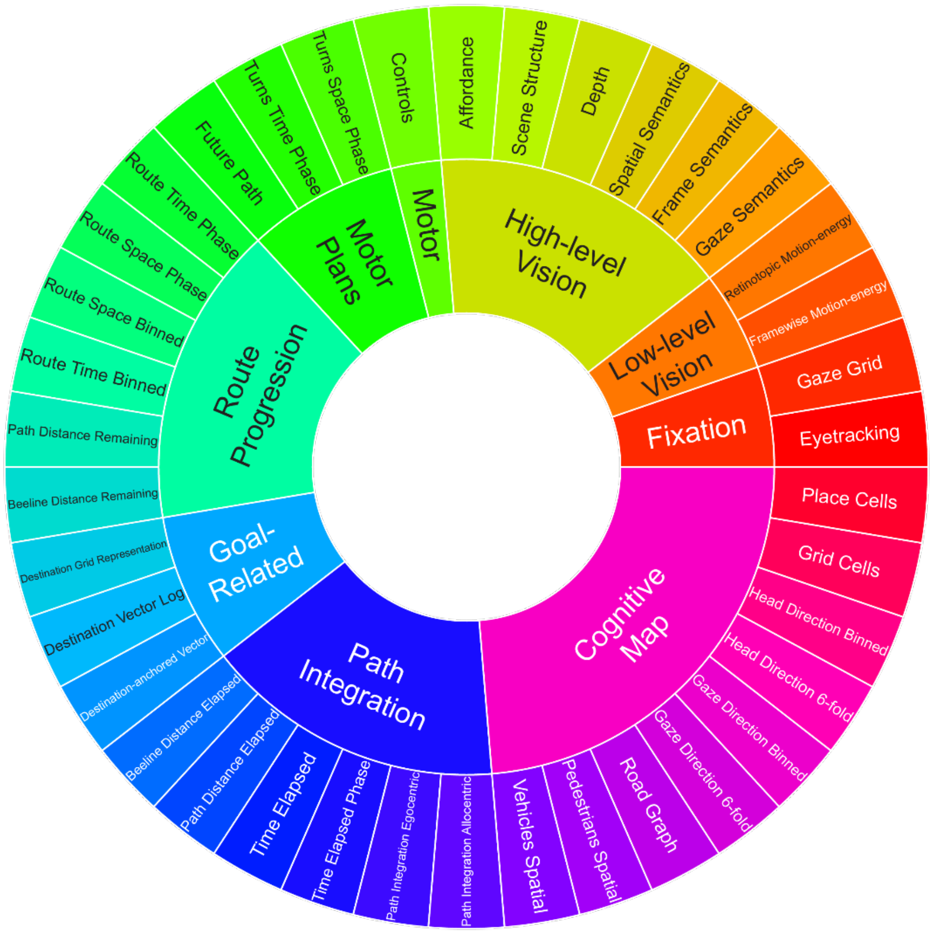


**Supplementary figure 6. Color legend for feature spaces and feature space categories in tuning bias profiles.**

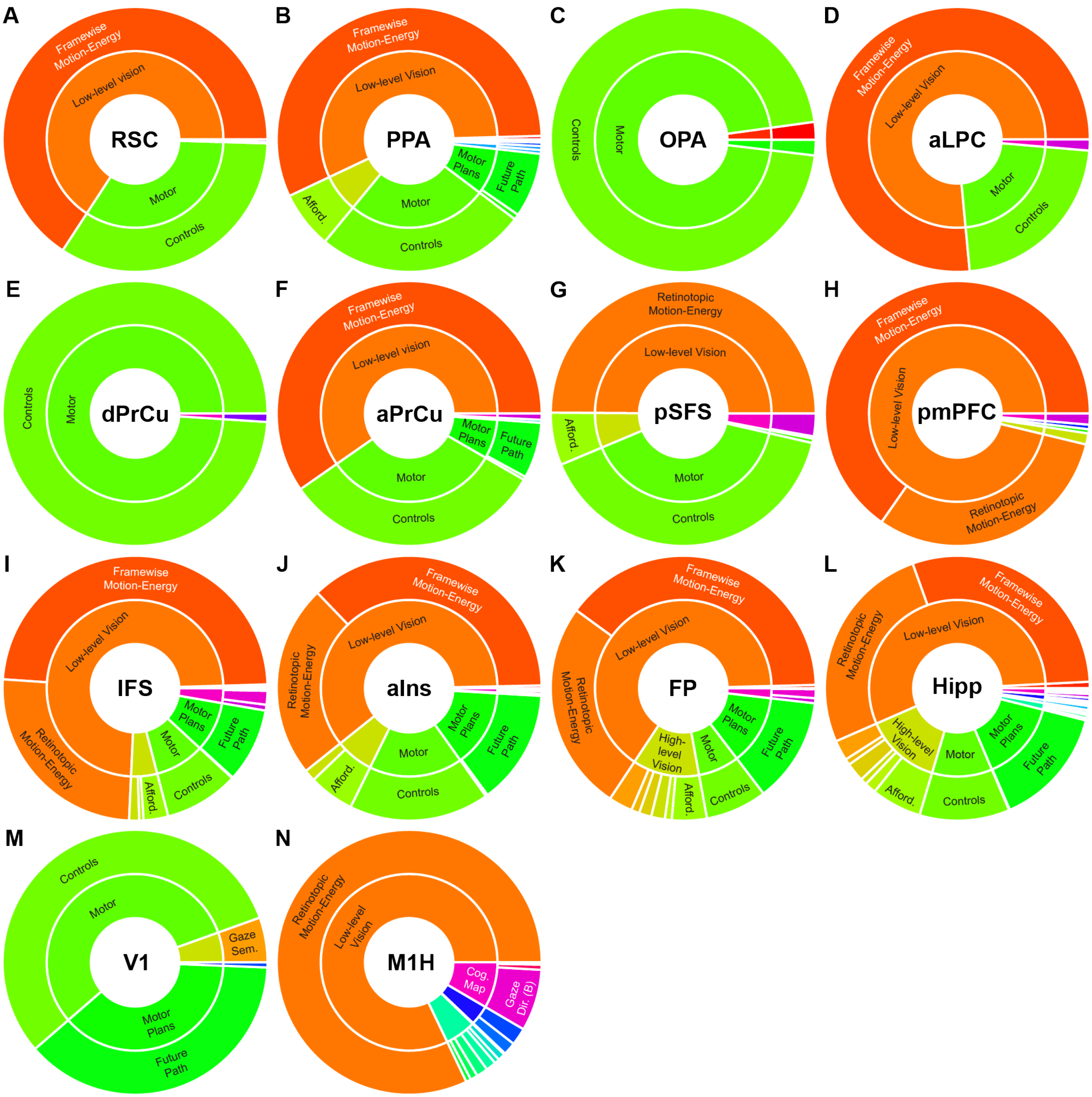


**Supplementary figure 7.** Feature spaces underrepresented by each of the ROIs in the navigation network relative to the rest of the cortex. These complement the overrepresented feature spaces by each ROI shown in Figure 3, and the figure follows the same format. A-L) The ROIs in the navigation network and the hippocampus primarily underrepresents low-level visual and motor control feature spaces. M) V1 underrepresents motor and motor plans feature spaces, and N) M1H underrepresents vision and navigation-related feature spaces.

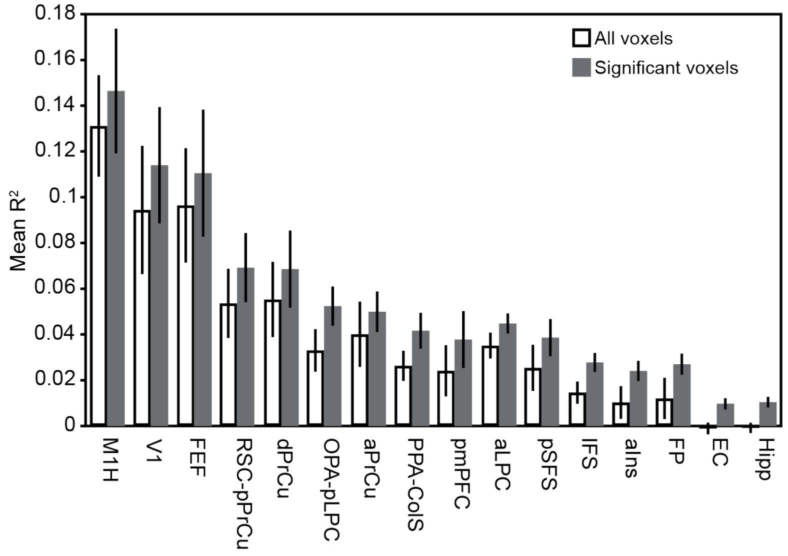


**Supplementary figure 8. SNR in medial temporal lobe structures.** Because of their physical distance from receiver coils and proximity to tissues that induce magnetic susceptibility artifacts, MTL regions exhibit much lower SNR than cortical regions. We quantified this difference by comparing the overall R2 in the entorhinal cortex (EC) and hippocampus with the 11 cortical regions identified in this study, and also the primary hand motor cortex (M1H), early visual cortex (V1), and frontal eye fields (FEF). White bars indicate average $R^2$ across all voxels in each ROI, and gray bars indicate average $R^2$ across significant voxels in each ROI. Error bars indicate standard deviation across subjects. Across the entire ROI, models explain no variance in the EC and hippocampus, and in significant voxels, models explain less than half the variance in the EC and hippocampus than in the least well-explained cortical region.

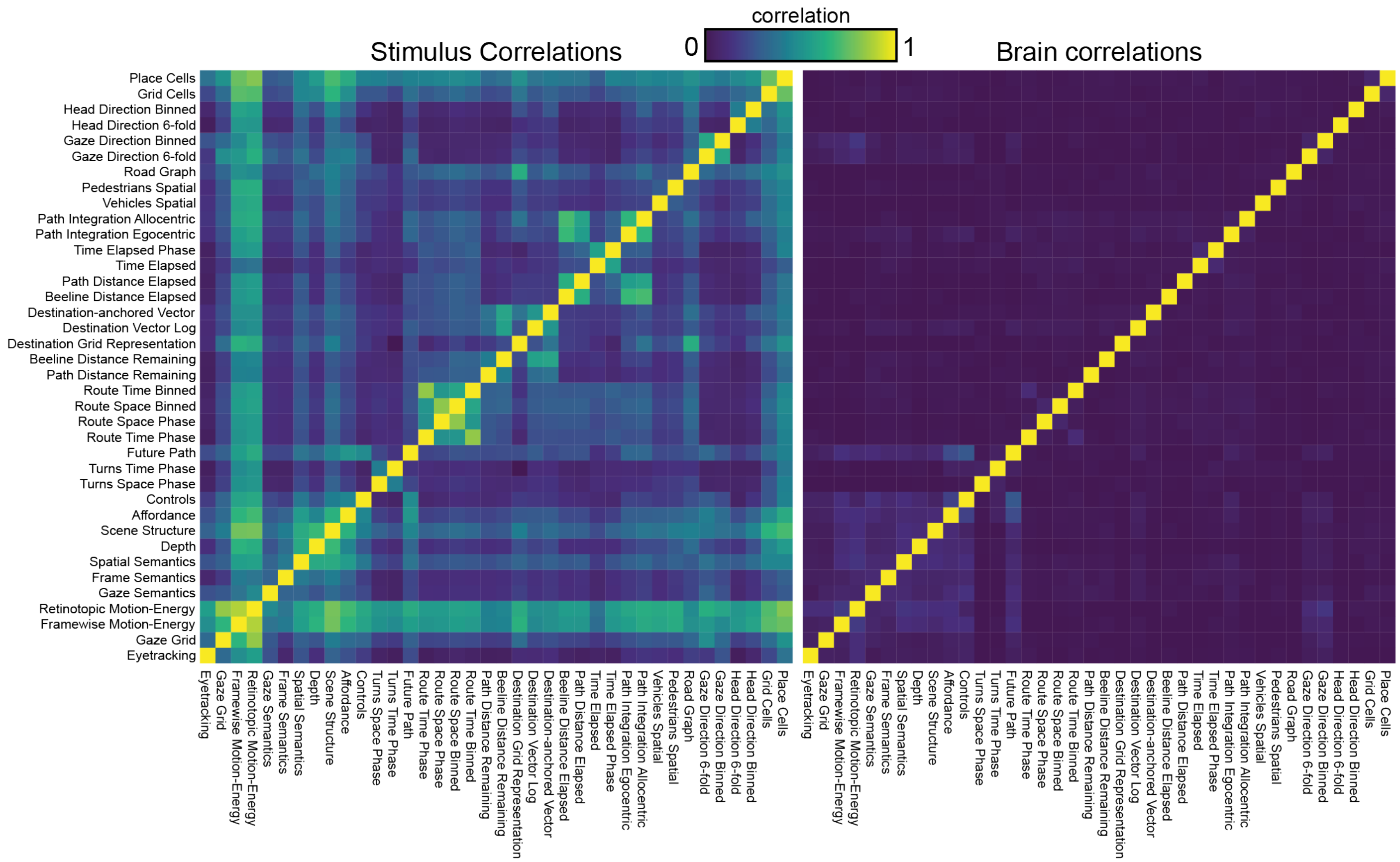


**Supplementary figure 9. Stimulus feature correlations and effects on model correlations.** A significant concern in naturalistic experiments is the inevitable correlations between features describing many different aspects of the task. To verify that the banded ridge models did not simply reflect correlations between feature spaces, we compared the correlations between feature spaces and the correlations between model performances for each feature spaces. The left panel shows correlations between feature spaces, while the right panels shows correlations between model performances for each feature space. Because feature spaces are of different sizes, the correlation between each pair of feature spaces was defined as the ordinary least squares average cross-prediction performance between the two feature spaces. Correlations between model performances was defined as the spatial correlation in prediction performance across the cortical surface for each pair of feature spaces. Results show that stimulus correlations do not explain any variance in the model correlations on the cortex ($R^2$ = -55.1±14.1, mean ± std.ev. across participants), indicating that the fit models are unlikely to be driven by stimulus correlations. These results demonstrate that banded ridge regression effectively disentangles correlated feature spaces during model fitting, enabling the analysis of data from complex naturalistic tasks.

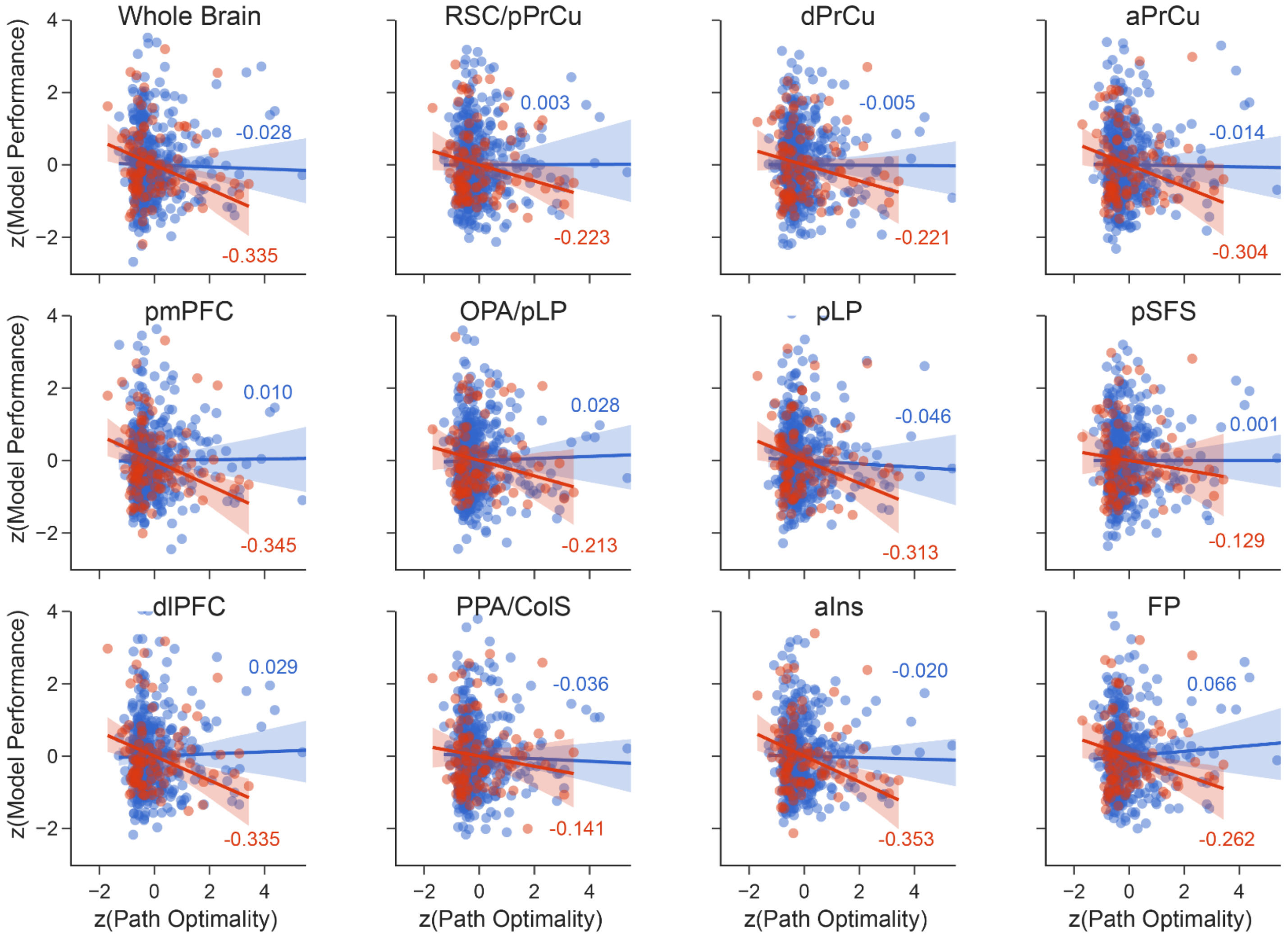


**Supplementary figure 10. Behavioral performance is weakly linked to encoding model performance**. Because the paths travelled by the participants were self-directed, behavioral variations should be reflected corresponding variations in brain activity. Thus, we sought to explore whether there are systematic relationships between model performance and behavioral performance To do so, we calculated model prediction performance in individual trials, and summed the split $R^2$ scores across models for navigation feature spaces. We then tested whether these navigation model performances were correlated with the path optimality on each trial, for both the whole brain and within each ROI in the cortical navigation network. To account for differences in SNR and performance across subjects, both the split $R^2$ scores and path optimality values are z-scored before aggregation. Because models perform better on the train data than test data, we separated trials from the train and test sets to avoid Simpson's paradox. In these plots, each dot is one trial, and blue indicates trials in train runs while red indicate trials in test runs. A trendline is shown along with a 99% confidence interval in the shaded areas. The correlation values are also shown for each distribution. While no relationship passed an FDR-corrected $p < 0.05$ significance threshold, these results suggest a general trend in decreasing model performance with more suboptimal paths in the test set. It is likely that because all subjects were performing close to ceiling, there is not enough variation in behavioral performance to establish significance. While these data hint that poor behavior may be linked to poorer representation of navigation-related information, future studies or analysis would be needed to better establish and characterize this relationship.

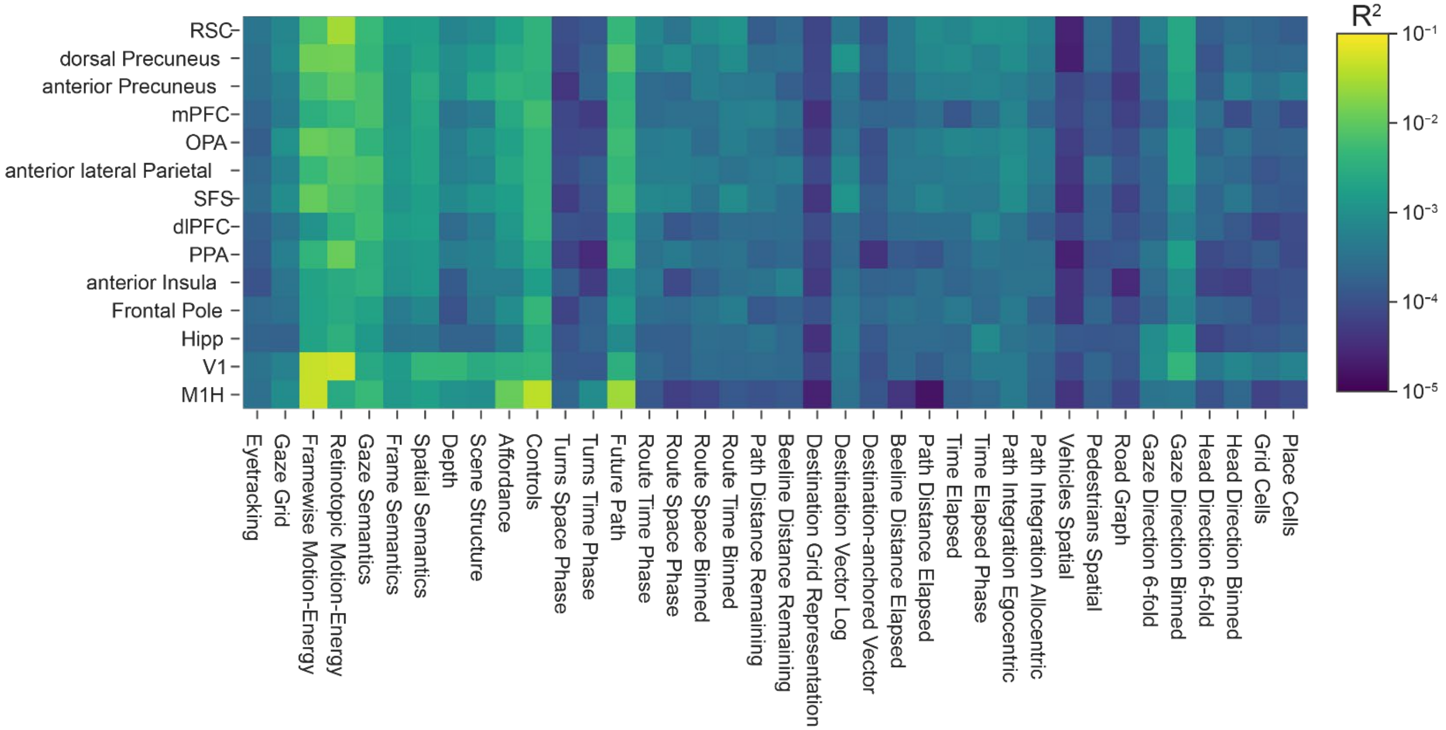


**Supplementary figure 11. Split $R^2$ scores for each of the navigation ROIs.** The average split $R^2$ scores for each feature space in each of the ROIs in the navigation network, the hippocampus, V1, and M1H. These values were used to determine the tuning bias profiles.

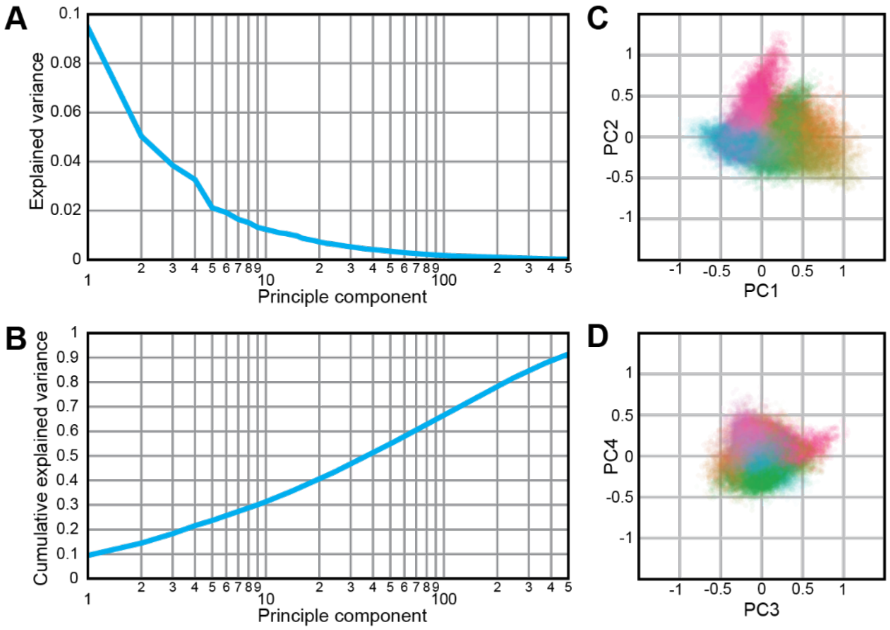


**Supplementary figure 12.** The encoding model weights are high-dimensional. A) The first PC explains less than 10 percent of variance in functional tuning in the cortex, and the fraction explained variance by the first 500 PCs of the weights of the navigation network follows a long-tailed distribution. B) The cumulative explained variance of the first 500 PCs. The first 3 PCs account for less than 20% of the variance in the weights, and 38 PCs are required to explain half the variance. Cortical vertices are projected to the planes formed by the 1st and 2nd (C) and 3$^{rd}$ and 4$^{th}$ (D) PCs, and each vertex is given the same color as it is in Fig. 4A. While the first two PCs show some correspondence to the organization in Fig. 4A, it is much less clear. The distribution is also notably non-Gaussian. The high-dimensionality and non-Gaussianity of the weights makes PCA not well-suited for describing the function organization of the cortex during navigation.

**Supplementary table 1.** List of feature spaces used for model fitting. For convenience of exposition, the 38 feature spaces are grouped into nine categories. A brief description and number of features are given for each feature space.

| Feature Space | # Features | Short Description |
|---|---|---|
| *Gaze-Related* | | |
| Eyetracking power | 1 | The aggregate power across channels encoding gaze location and its derivatives in screen space encoded in cartesian and polar coordinates |
| Gaze grid | 288 | Gaze location in screen space encoded with a hexagonal grid basis [1] |
| *Low-level vision* | | |
| Framewise motion-energy | 8418 | Spatiotemporal Gabor energy in screen coordinates [2] |
| Retinotopic motion-energy | 8418 | Spatiotemporal Gabor energy in retinotopic coordinates |
| *High-level vision* | | |
| Frame semantics | 33 | Semantic content of the screen |
| Gaze Semantics | 33 | Semantic content of a 5° circle around the gaze location |
| Depth | 30 | Distance to objects in the virtual world |
| Scene structure | 637 | The distribution of visible surfaces and their orientations relative to the participant's viewpoint [3] |
| Spatial semantics | 672 | Visible semantic content of the virtual world around the participant in egocentric coordinates |
| Navigational affordance | 160 | Where in the scene that the participant can move through [4] |
| *Motor* | | |
| Controls | 4 | Steering, gas, brake, and gear selections from the subject |
| *Motor plans* | | |
| Future path | 160 | Future path that the participant will take in egocentric coordinates |
| Distance to next turn | 24 | Fraction of total distance travelled until the next turn encoded with a Fourier basis [5] |
| Time to next turn | 24 | Fraction of total time travelled until the next turn encoded with a Fourier basis |
| *Route progression* | | |
| Beeline distance remaining | 12 | Beeline distance remaining to the destination [6] |
| Path distance remaining | 12 | Distance remaining along the actual planned path to the destination [6] |
| Spatial route progression (continuous) | 8 | Fraction of total distance travelled to the destination encoded by a Fourier basis [7] |
| Temporal route progression (continuous) | 8 | Fraction of total time travelled to the destination encoded by a Fourier basis |
| Spatial route progression (binned) | 43 | Fraction of total distance travelled to the destination encoded by a set of indicator features |
| Temporal route progression (binned) | 43 | Fraction of total time to the destination encoded by a set of indicator features |
| *Goal-related* | | |
| Destination vector | 97 | An egocentric vector to the destination [6,8,9] |
| Destination-anchored vector | 97 | An allocentric vector from the destination to the current location of the participant |
| Destination grid representation | 337 | Position of the destination in the world encoded with a hexagonal grid basis |
| *Path Integration* | | |
| Allocentric path integration | 97 | Allocentric vector to the start location of the current trial [10,11] |
| Egocentric path integration | 97 | Egocentric vector to the start location of the current trial |
| Beeline distance elapsed | 12 | Beeline distance from the start of the current trial [10,11] |
| Path distance elapsed | 12 | Actual path distance travelled from the start of the current trial |
| Time elapsed | 50 | Time elapsed since the start of the current trial encoded by a set of indicator features |
| Time elapsed phase | 10 | Time elapsed since the start of the current trial encoded by a Fourier basis |
| *Cognitive map* | | |

| | | |
|---|---|---|
| Place fields | 6579 | Place field-like representations of the participant's 2D location in the virtual world[12,13] |
| Grid cells | 336 | Features with grid cell-like periodic spatial selectivity on a hexagonal grid in the virtual world [14,15] |
| Head direction | 8 | The head direction of the participant in the world encoded with a set of indicator bins [16,17] |
| 6-fold Head direction | 2 | The head direction of the participant in the world, with 6-fold symmetry, encoded with a quadrature filter [18] |
| Gaze direction | 8 | The gaze direction of the participant in the world encoded with a set of indicator bins |
| 6-fold gaze direction | 2 | The gaze direction of the participant in the world, with 6-fold symmetry, encoded with a quadrature filter |
| Road graph | 1042 | The planned path of the subject expressed as a set of edges on a graphical representation of the road network [19–21] |
| Pedestrians | 160 | Positions of pedestrians in egocentric coordinates [22,23] |
| Vehicles | 160 | Positions of other cars in egocentric coordinates |

### Supplementary Methods

Localizers for known ROIs

Conventional functional ROIs were identified in each participant in separate functional localizer experiments. ROIs were determined using five sets of localizers: a retinotopic localizer, a MT localizer, a visual category localizer, a motor localizer, and a theory-of-mind localizer.

*Retinotopic Localizer*

Retinotopic localizers were collected in four 9-minute runs. In two runs, rotating wedges were used to determine visual angles. In the other two runs, expanding rings were used to determine eccentricity. Visual angle and eccentricity maps were then used to delineate V1, V2, V3, V4, V3A, V3B, and V7 [24].

*MT Localizer*

MT localizer data were collected in four 90-second runs. These runs consisted of alternating 16-second blocks of motion coherent and incoherent random dot fields. The contrast between coherent and incoherent blocks was used to delineate MT+.

*Visual category localizer*

Visual category localizer data were collected in six 4.5-minute runs. These runs consisted of 16 blocks of 16 seconds each. In each block, 20 images of either places, faces, body parts, non-human animals, household objects, or phase-scrambled objects were displayed. Each image was displayed for 300 ms followed by a 500 ms blank. The contrast between faces and objects was used to define the fusiform face area (FFA, [25]). The contrast between body parts and objects was used to define the extrastriate body area (EBA, [26]). The contrast between places and objects was used to define the parahippocampal place area (PPA, [27]), occipital place area (OPA, [28]), and retrosplenial cortex (RSC, [29]).

*Motor localizer*

Motor localizer data were collected in one 10-minute run. Participants were cued to randomly alternate between six motor tasks in 20-second blocks. For "hand," "foot," "mouth," "speech," and "rest," the cue was simply that word presented at the center of the screen. For the saccade condition, participants were shown a screen filled with randomly distributed saccade targets.

In the "hand" blocks, participants were instructed to make small finger-drumming movements. In the "foot" blocks, participants were instructed to make small toe movements. In the "mouth" blocks, participants were instructed to make small lip movements without moving the tongue or jaw. In the "speak" blocks, participants were instructed to continuously subvocalize self-generated sentences without physically speaking. In the "saccade" blocks, participants were instructed to continuously saccade between targets presented on the screen. A linear model was used to find the change in BOLD response in each voxel in each condition relative to the mean BOLD response.

Weights for the hand, foot, and mouth responses were used to delineate the primary motor and somatosensory areas for hands (M1H, S1H), feet (M1F, S1F), and mouth (M1M, S1M). Weights for the saccade responses were used to delineate the intraparietal sulcus (IPS), frontal eye fields (FEF, [30]), and frontal operculum (FO [31]). Weights for the speak responses were used to delineate Broca's area and the superior ventral premotor speech area (sPMv).

*Theory-of-mind localizer*

Theory of mind localizer data were collected in two 5-minute runs. Participants were shown a series of short stories consisting of false belief, false photograph, desire, physical description, and nonhuman description stories [32]. Each story was shown for 10 seconds. After each story, participants were presented with a fill-in-the-blank two-alternative forced choice question for 4 seconds and responded with a buttonbox. Half of the questions concerned the belief of characters in the stories, while the other half were concerned with factual information. The contrast between belief and factual questions was used to delineate the temporoparietal junction (TPJ).

<u>Relationship between voxelwise encoding models and general linear models</u>

As noted in the main text, both the voxelwise encoding modelling (VEM) framework and the older statistical parametric mapping (SPM) [33] framework in human neuroimaging build general linear models (GLMs) of the form

$$Y = XW + \epsilon$$

here brain activity Y is modelled as a linear combination of regressors X, the weights W are estimated during model fitting, and ε is noise. However, while both the VEM and SPM frameworks make use of a GLM, the two frameworks differ in experimental design, model fitting, and interpretation of models. Here we describe the differences in the VEM and SPM frameworks for solving the GLM problem. For clarity, we use "GLM" to refer to a mathematical model of the form $Y = XW + \epsilon$, and "VEM" and "SPM" to refer to the integrated neuroimaging experimental frameworks that both solve a mathematical GLM problem.

First, the SPM and VEM frameworks take different approaches to experimental design. Because SPM typically relies on ordinary least squares (OLS) regression, it requires structured experimental designs in which explicitly defined conditions or events are presented to participants. Such experiments are also usually designed to minimize correlations between experimental variables. The operationally-defined conditions or events of interest are then modelled as separate regressors in X. This approach typically assumes that the conditions and event regressors can be designed to be orthogonal. The regressor matrix X then typically consists of relatively few columns that capture temporally orthogonal experimental conditions. In contrast, because VEM uses robust model estimation techniques, it can be applied to naturalistic experiments that do not have clear conditions or events. In order to preserve the statistical structure of naturalistic experiments, correlations between stimulus features are typically not removed. Time-varying stimulus and task features, such as semantic embeddings or image filters, are used as regressors in X, and correlations between features are addressed in the model fitting process. The VEM approach makes no assumption about the orthogonality of regressors. In many experiments that use VEM, the regressor matrix X is large, reflecting many hundreds or thousands of time-varying stimulus features. In many cases, such as in the experiment presented here, the regressor matrix X may have more columns than time samples.

Second, the SPM and VEM frameworks use different model fitting processes for estimating W. The SPM framework uses ordinary least squares (OLS) regression to estimate W and to perform statistical inference on the parameters. However, OLS requires more time samples than regressors, and may produce biased estimates when regressors are collinear. Furthermore, by solely minimizing error on the training data, OLS overfits and generalizes poorly to new data. These properties make the SPM framework poorly-suited for modelling data from more naturalistic experiments. In contrast, the VEM framework uses regularized regression to estimate W, and thus avoids many of the pitfalls of OLS. By penalizing large model weights, regularized regression more effectively accounts for collinear regressors by minimizing their weights. The shrinkage in weights also reduces overfitting and improves generalization to new data. Finally, regularized regression also enables models to be fit even when there are more regressors than time samples. Early versions of VEM used ridge regression to apply a uniform regularizer to all regressors. The current version of VEM uses banded ridge regression, which allows groups of regressors to be regularized separately. The VEM framework places heavy emphasis on cross-validation and out-of-set predictions to guard against overfitting, which avoids overfitting to training data or the regularization hyperparameters, and ensures generalization to new data. Model quality is explicitly evaluated by prediction accuracy on out-of-set data.

Finally, the SPM and VEM frameworks differ in how the fit models are interpreted. The SPM framework focuses on hypothesis-driven contrasts, classical statistical inference, and significance testing of parameter estimates in the weight matrix W. Contrasts and statistical significance are used to identify brain regions that respond more to one experimentally-defined condition or event than another. In contrast, the VEM approach emphasizes the representational content of brain activity. The weight matrix W is used to predict brain activity in response to new inputs, and also to provide detailed descriptions of what stimulus and task features drive brain responses. Because both approaches use a general linear model, the weights W produced in VEM can also be used for statistical inference and computing contrasts. However, in practice, naturalistic stimuli and tasks rarely sample experimenter-defined contrasts and conditions in the same manner as controlled stimuli from SPM experiments.

In sum, while both the VEM and SPM frameworks solve some form of a mathematical GLM problem, the VEM framework offers greater flexibility and suitability for the current experimental setting. Unlike the SPM approach, which relies on structured designs with orthogonal regressors, the VEM framework accommodates high-dimensional, collinear regressors through regularization, cross-validation, and prediction. In continuous, naturalistic settings without clearly defined experimental conditions, hypothesis-driven contrasts become less meaningful and hinder the applicability of the SPM framework. The flexibility of the VEM framework is a more appropriate and powerful method for understanding brain responses in these contexts.

### Feature space descriptions

To capture the many different types of information that may be encoded by the brain during active, naturalistic navigation, we developed 38 feature spaces that capture a variety of perceptual-, motor-, and navigation-related features. These feature spaces were drawn from both the existing human and animal navigation literature, and also reflect features that we hypothesized the brain to represent. For convenience, we grouped these 38 feature spaces into nine categories: two fixation-related feature spaces that capture eye movement-related information, two low-level vision feature spaces that capture the 2D structure of the stimulus in screen space, six high-level vision feature spaces that capture 3D and semantic structure of the stimulus in virtual world space, one motor feature space that capture the concrete motor actions of participants, three motor plans feature spaces that capture planned actions, six route progression feature spaces that capture progression towards the destination, three goal-related feature spaces that capture information about the location of the destination, six path integration feature spaces that capture the participant's progress from the start location on every trial, and nine cognitive map feature spaces that capture information represented in an allocentric world map.

Here we describe how features are computed for every feature space. Unless otherwise noted, the features were computed at 15 frames per second (fps) and then integrated across each TR.

### Gaze-related feature spaces

#### *Eyetracking power*

A set of eyetracking features were included to capture effects related to eye movements that are not directly reflected by the retinotopic motion-energy features. The eyetracking data was used to determine (x, y) gaze position in screen coordinates. The origin of these coordinates was set to the center of the screen. For each dimension (x and y), the second and third powers and the first, second, and third derivatives of the position values were also computed. The conversion of the cartesian coordinates into polar coordinates was also included. Finally, each channel was rectified, and we computed the average power across all channels to produce 1 feature at 60 fps.

*Gaze grid*

Non-human primate electrophysiology suggests that the gaze position in a head-centered reference frame is also represented on a hexagonal lattice basis [34]. There has also been some fMRI evidence for such representations in the human brain [1]. We therefore built a set of features to represent gaze location using a grid cell-like hexagonal basis. We used hexagonal lattices with spacings of 16, 32, 64, 128, 256, and 512 pixels in screen space. For each lattice, the lattice was constructed by placing 2D Gaussian functions at the vertices. These Gaussian functions had a standard deviation of 1/12 the spacing of the lattice to account for the phase-offset copies of the lattice. Each lattice forms a basis function. For each spacing, we included variants with spatial phase offsets of 1/3 and 2/3 the spacing, and also rotated copies at 20 and 40 degrees of rotation. At each frame, the value of each basis function was evaluated at the gaze position in screen space for a total of 162 features.

<u>Low-level vision feature spaces</u>

*Framewise Motion-Energy*

The motion-energy feature space consists of a pyramid of spatiotemporal Gabor filters that captures the low-level visual structure of the stimulus [2]. First, screen recording videos were downsampled from 1024 × 768 to 128 × 96 pixels. This 128 × 96 pixel image was centered on a uniform gray (12.5% brightness) background to match the frame size of the retinotopic motion-energy features (see next section). Next, the color frames were converted to grayscale. The spatiotemporal Gabor pyramid was created using 7 spatial frequencies (0, 1, 2, 4, 8, 16, and 32 cycles across the image), 3 temporal frequencies (0, 2, and 4 Hz), and 8 orientations (0, 45, 90, 135, 180, 225, 270, and 315 degrees). These filters tiled the screen, and the video frames were convolved with these filters. Finally, filter responses were squared and summed for each quadrature pair for a total of 8418 features. These features were computed at 30 fps.

*Retinotopic Motion-Energy*

Because participants were freely viewing the screen, saccades across the stimulus will produce large changes in the images seen by the retina. To account for these effects, the motion-energy filter responses were also calculated in retinotopic coordinates rather than in screen coordinates. First, the eyetracking data was used to determine the gaze location for each frame of the stimulus. Second, the stimulus videos were downsampled to from 1024 × 768 to 128 × 96 pixels, and the gaze location on each frame was also rescaled to match this downsampled size. Then, each downsampled frame was placed into a 256 × 192 uniform gray (12.5% brightness) background such that the gaze position is centered in this larger frame. Finally, motion-energy features were computed using these frames with the same methods for computing frame-wise motion-energy features for a total of 8418 features. These features were computed at 30 fps.

<u>High-level vision feature spaces</u>

*Frame semantics*

The frame semantics feature space captures the visual semantic content of the entire screen. To compute the frame semantic features, we used the game engine to apply semantic segmentation to the participant's view to determine the category of the object rendered at each pixel. We used 16 categories that were relevant to driving: buildings, pedestrians, vehicles, roads, road lines, sidewalks, traffic signs, poles, walls, fences, fields, ground, foliage, self, miscellaneous objects, and sky. We also included features for on-screen display items that were overlaid on the screen rather than rendered in-world; these are the "go to" and "arrived" cues and the speedometer. This feature space also included 14 additional categories for objects and on-screen display items that appeared during the learning phase, but not during data collection. The feature values were calculated as the fraction of the frame occupied by each category. This feature space contains a total of 33 features.

*Gaze semantics*

Participants are unlikely to allocate equal attention to all areas of the screen. Thus, the frame semantics feature space may not accurately reflect representations of visual semantics with the influence of top-down attention. Because participants were freely viewing the screen, we used the gaze location as a proxy of how participants allocated attention. We therefore created a set of visual semantic features for the part of the screen at which the participant is fixating. The semantic segmentation of the scene from the frame semantics feature space was re-used here. For each time sample in the eyetracking data, a circle of 5° diameter was drawn around the gaze position on the corresponding semantic segmentation frame. Then the fraction of this circle that was occupied by each category was calculated, producing 33 features. These features were computed at 60 fps.

*Depth*

The depth feature space captures the distribution of distances to visible objects in the virtual world. To compute the depth features, we used the game engine to determine the distance to the surface rendered at each pixel. Distances are encoded up to 1000 m with a precision of 0.06 mm. Then, the frame is divided up into 5 horizontal bins that each span 22 degrees in the virtual environment. Six distance bins at 0-5 m, 5-10 m, 10-25 m, 25-50 m, 50-100 m, and 100+ m were created. In each horizontal bin, we then counted the number of pixels that fell within each distance bin for a total of 30 features.

*Scene structure*

To capture the geometry of the scene, we created features that described the surfaces, their orientations in the frame, and relative position to the participant [3]. These features utilized the pixel-wise depth information calculated for the depth feature space. We used the game engine to also compute the normal vector (relative to the camera) of the surface at each pixel. We divided the frame up into 7 horizontal bins that each span 15.7 degrees in the virtual environment. We then created 10 log-spaced distance bins between 0 and 1000 m from the participant. We then created 9 surface orientation bins: up, down, left, right, straight ahead, and four oblique directions. Within each combination of horizontal and distance bin, pixels were sorted into the closest of the 9 orientation bins. Additionally, at each horizontal bin, we included one extra channel for the sky, which has an undefined orientation vector. This feature space contains a total of 637 features.

*Spatial semantics*

Both the frame and gaze visual semantic feature spaces are computed in the 2-dimensional image space of the eye or screen. To be navigationally useful, these semantics must be converted into the 3-dimensional physical space of the world. We built the spatial semantic feature space to capture the 3D spatial distribution of visual semantics around the participant in the virtual world. First, we combined the depth and semantic segmentation information to create a combined distance-and-object-type label for each pixel. Here, we excluded the on-screen display items because they are not in the physical space of the virtual world, and used only the 16 object categories for in-world objects. Then, we divided the frame up into 7 horizontal bins, each spanning 15.7 degrees in the virtual environment, and 6 distance bins at 0-5 m, 5-10 m, 10-25 m, 25-50 m, 50-100 m, and 100+ m. Finally, for each combination of horizontal and distance bins, we counted the number of pixels in each of the 16 in-world semantic categories in that bin for a total of 672 features.

*Navigational affordances*

Navigational affordances have been suggested to be represented in the OPA [4,35]. Here, we define navigational affordances as where the participant is legally allowed to drive. As such, this feature space essentially captures the roads and parking lots around the participant. We use the demo files to parse the location and heading of the participant's car at each frame. We then build a series of log-polar spatial bins around the participants' position. There are 16 directions, and in each direction, there are 10 log-spaced distance bins from 0 to 100 m away from the participant. The bins are rotated such that a reference direction is always aligned with the participant's heading at each frame. We term this bin layout "egocentric log-polar bins" to reflect that they are referenced to the immediate heading of the participant, that the distance bins are log-spaced, and that they are in polar coordinates.

We constructed a map of the extent of all the roads and parking lots in the map. Each bin is then compared against this map, and the fraction of each bin that overlaps with this navigable space is recorded as the feature value. Unlike the previous feature spaces, the affordance features are not computed from the video frames, despite the semantic segmentation providing a trivial method of finding roads in the video frames. We do so to capture affordances that are beyond the current visible space, as the view in the virtual world can only show what is in front of the participant. This feature space contains a total of 160 features.

Motor feature spaces

*Controls*

The motor actions of the participants are directly reflected by the corresponding control inputs. We therefore included features for steering wheel angle, brake input, accelerator input, and gear selection. The steering wheel angle, brake input, and accelerator inputs were continuous values, while the gear selection (toggle between forward and reverse) was a binary indicator for a button press. This feature space contains a total of 4 features.

Motor plans feature spaces

*Future Path*

A plan is needed for successful navigation. We operationally divide the plan into two scales: the long-term abstract plan and the short-term concrete plan. The future path feature space captures the short-term concrete plan. This feature space metrically represents where the participant plans to move in the visible world and the immediately adjacent non-visible space. We again make use of egocentric polar bins. In each of 16 directions, we place 10 bins log-spaced from 2.5 m to 100 m away from the participant. At each frame, we use the demo files to look ahead in time for the locations of the participant in future frames. We search forwards in time to either when the participant is beyond the furthest edge of the bins (100 meters from the current location), or until the end of the current navigational segment, whichever is earlier. Then, for each bin that the participant will be in in the future, we set its value to 1. We set all other bins values to 0. These 160 features capture the near-term navigational plan of where the participant plans to go.

*Distance to next turn*

A route can be divided into smaller linear segments joined by turns. Rodent neurophysiology has shown that PPC neurons are selective for these individual segments [5]. To capture any such representations in our experiment, we first define "linear segments" as segments between turns at intersections. Each navigational segment was manually broken up into linear segments, separated by turns. We then built pairs of sinusoid functions with 90 degree phase offsets that swept through 1, 2, 4, and 8 cycles along the length of each linear segment. These phase-offset features allow us to capture selectivity to any fractional position along linear segments. At each frame, the position of the participant along the linear segment was used to calculate the value of these sinusoids. Because left turns, right turns, and U-turns may have distinct representations, we created a separate set of features for each type of turn. At any given point, only features for the particular upcoming turn type were populated, and the features for the other two turn types were zeroed. Because the sinusoidal functions were duplicated across three turn types, there are a total of 24 features.

*Time to next turn*

We also created a set of features that encoded the time until the next turn. While the time and distance remaining are correlated, the correlation is perfect. Thus, we also built a temporal version of the turn progression features. We determined the time it took participants to drive through each linear segment, and built pairs of sinusoid functions with 90 degree phase offsets that swept through 1, 2, 4, and 8 cycles for each segment. At each frame, the time elapsed since the beginning of the current linear segment was used to compute the value of these functions. We again created duplicates of these sinusoids for the three types of possible turns for a total of 24 features.

<u>Route progression feature spaces</u>

*Beeline distance remaining*

Human neuroimaging experiments have suggested that the hippocampus and entorhinal cortex may encode both the path and beeline distance to goals [6]. At every frame, we therefore computed the beeline distance between the participant and the destination. Like the beeline distance elapsed features, the log transform of the beeline distance remaining was soft-binned into 12 bins equally spaced between 1 and 12 log units for a total of 12 features.

*Path distance remaining*

To complement the beeline distance remaining feature space, we also computed the path distance remaining [6] between the participant and the destination. At every frame, we used game recordings to look ahead in time, and calculated the distance that the participant will travel from their current position to the destination. The log transform of this distance was also soft-binned into 12 bins equally spaced between 1 and 12 log units for a total of 12 features.

*Spatial route progression (continuous)*

The route progression feature spaces were inspired by a rodent study that found RSC neurons that demonstrated spatially periodic firing fields when animals ran along a track [7]. Some neurons had a single firing field, while others showed 2 or 4 firing fields evenly spaced along the track. These firing field periodicities were conserved across differently-shaped tracks, suggesting that they encoded generalized position along any path. This study modeled the firing fields with pairs of sinusoidal functions that swept through 1, 2, 3, 4, 6, and 8 cycles along the length of the track. The sinusoids are offset in phase by 90 degrees, and form quadrature filters that can capture periodic activity without any assumptions about phase.

Here, we adapted these features to the paths that the participants took in the virtual world. On each trial, the path that the participant took. Each route was then normalized to a length of 1. Then, we built pairs of sinusoid functions with 90 degree phase offsets that swept through 1, 2, 4, and 8 cycles along this linearized path. At each frame, we used the position of the participant along the path to compute the value of these functions for a total of 8 features.

*Temporal route progression (continuous)*

The temporal route progression feature space is closely related to the spatial route progression feature space. While the spatial route progression features were based on the distance elapsed, the temporal progression features are based on time elapsed. Unlike the spatial progression features, these temporal progression features are not speed-dependent. For example, when the participant is waiting at a red light, the spatial route progression features remain constant, while the temporal progression features continue to change.

For each trial, we determined the total time that the participant took. Then, we built pairs of sinusoid functions with 90 degree phase offsets that swept through 1, 2, 4, and 8 cycles in this amount of time. At each frame, we used the time elapsed since the beginning of the segment to compute the value of these functions for a total of 8 features.

*Spatial route progression (binned)*

The Fourier encoding of route progression implicitly assumes that neurons display periodic tuning to progression along the route. In [7], some cells were shown to be partially periodic, e.g. they displayed spatially periodic firing fields in one half of the route. Such tuning profiles cannot be easily captured by the Fourier basis features in a linear model, as they require interactions between features. Thus, we created a complementary set of features, which divide the route into 1, 2, 4, 8, 12, and 16 bins. The 1-bin feature serves as an indicator that the participant is actively navigating to a destination, rather than waiting during the intertrial delay period. At each frame, we determined which spatial portion of the route that the participant is in, and set the corresponding feature values to 1 and others to 0 in each set of bins. This feature space contains a total of 43 features.

*Temporal route progression (binned)*

We applied the same rationale to the temporal route progression feature space. We created a corresponding set of binned temporal progression features that divide the route into 1, 2, 4, 8, 12, and 16 bins. At each frame, we determined which temporal portion of the route that the participant is in, and set the corresponding feature values to 1 and others to 0 in each set of bins. This feature space contains a total of 43 features.

Goal-related feature spaces

*Destination vector*

Previous studies have shown that the human brain tracks proximity to a goal [6,8] and that rodent hippocampal place cells encode goal vectors [9]. We therefore built a feature space to encapsulate an egocentric vector to the destination. At every frame, we determined the position and heading of the participant, and also the location of the destination. From these values, we determined an egocentric vector from the participant to the destination in polar coordinates. We then took the log transform of the distance component of this vector. In polar egocentric space, we constructed a set of 2D Gaussian functions. In each of 16 equally-spaced radial directions, we placed the center of 12 equally-spaced 2D Gaussians functions between 1 and 12 log units from the participant. The standard deviation of these Gaussian functions was set to be 1/6 that of the distance between neighboring bins. The value of these Gaussian functions at the log-transformed destination vector were used as features. We term this process *soft-binning* the values. Features were zeroed during the periods between successive navigational segments. We include an additional indicator variable that is 0 when the participant is navigating, and 1 during the intertrial delay period. This feature space contains a total of 97 features.

*Destination-anchored vector*

The destination vector is an egocentric vector anchored to the current position and heading of the participant. An equally valid description of the relationship between the participant and destination could be anchored to the destination. We therefore also took the log-distance allocentric vector from the destination to the position of the participant, and soft-binned this vector at every frame into another set of 97 features. Note that these features are not equivalent to the destination vector. Whereas destination vector features overrepresent the space around the participant, the destination-anchored vector over-represents the space around the destination.

*Destination grid representation*

Place cell spatial firing fields can be derived from multiple grid cell inputs[36]. Thus, at any moment, the participant's position can be encoded using only the grid cell features. Because the destination is of particular interest during navigation, we postulated that its position may also be represented. Therefore, we created a second set of grid cell features with spacings of 480 m, 240 m, 120 m, 80 m, 40 m, 20, and 10 m in physical space. For each spacing, we included variants with spatial phase offsets of ¼, ½, and ¾ the spacing, and also rotated copies at 20 and 40 degrees of rotation. During each trial, these features were used to represent the position of the destination. We included one indicator variable for the intertrial period when there is no destination. This feature space contained a total of 190 features.

<u>Path integration feature spaces</u>

*Allocentric path integration*

Neuroimaging studies suggest that both the hippocampus and RSC may represent path integration features [10,11]. To capture this information, on each trial, we first defined the starting position as the position of the participant during the "go to" cue prompt. We then computed a homing vector referenced to the global north. The distance component is log-transformed, and the resulting vector is soft-binned into 8 directions with 12 distances per direction. We include an indicator variable that is 0 when the participant is navigating, and 1 during the intertrial delay. This feature space contains a total of 97 features.

*Egocentric path integration*

The path integration vectors in the allocentric path integration feature space is referenced to the global north. To capture any egocentric representations, we computed a second set of path integration vectors referenced to the current heading of the participant. Aside from the reference direction, these features were computed in the same manner as the allocentric path integration features. This feature space contains a total of 97 features.

*Beeline distance elapsed*

Path integration is commonly assumed to be computed with respect to some starting position, and neuroimaging studies suggest that both the hippocampus and RSC may represent path integration features [10,11]. For each navigation segment, we defined the starting position as the position of the participant during the “go to” cue prompt. At each frame, we computed the beeline distance from the current position of the participant to the starting position. The log transform of this distance is then soft-binned into 12 bins equally spaced between 1 and 12 log units for a total of 12 features. Like the destination vector features, the feature values were zeroed in the periods between successive navigational segments. Note this feature space is not equivalent to the beeline distance remaining feature space: the log spacing overrepresents distances close to the starting position in this feature space, rather than distances close to the destination.

*Path distance elapsed*

Very rarely do participants travel in a straight line between two points. It is thus possible that, in addition to the beeline distance, the brain also tracks the actual distance traveled. At every frame, we computed the actual distance traveled from the starting position. The log transform of this distance is then soft-binned into 12 bins equally spaced between 1 and 12 log units for a total of 12 features. Note that unlike the route progression features, the path distance elapsed features are not normalized, and metrically reflect the distance traveled.

*Time elapsed*

In addition to tracking the distance elapsed, the brain may also track the time elapsed since the beginning of the trial. While time elapsed is correlated with distance travelled, it is not perfectly correlated. For example, when the participant is waiting at a red light, time elapsed continues to increase, but distance elapsed does not. Elapsed time was captured with a set of binned features that represented time elapsed in 10-second intervals ranging from 10 to 500 seconds. This feature space contains a total of 50 features.

*Time elapsed phase*

The time elapsed feature space uses a set of indicator features that each uniquely represent an amount of elapsed time. However, this parameterization is not well-suited for capturing any periodic representation of time elapsed. Thus, we also created a set of periodic features for the elapsed time. These features were implemented as pairs of sinusoids with a 90 degree phase offset, with periods of 15, 30, 60, 90, and 120 seconds. This feature space contains a total of 10 features.

<u>Cognitive map feature spaces</u>

*Place Fields*

Both animal and human electrophysiology have demonstrated the existence of place cells [12,13]. We therefore included place field features that tiled the map at 480 m, 240m, 120m, 80m, 40m, and 20 m intervals. Each place field is modeled as a spatial Gaussian field with a standard deviation of 1/5 of the tiling distance. Across all spacings, there are a total of 14,144 possible fields. Because these fields tile the map, many are in inaccessible locations, such as the center of a lake or inside buildings. Additionally, some fields lie on side roads that participants did not use. Therefore, we pruned the place fields down to the union of the sets of place fields traversed by each participant. This feature space contains a total of 6579 features.

*Grid cells*

Rodent electrophysiology studies showed that cells in the entorhinal cortex metrically measure space using a hexagonal lattice basis [14,15]. We therefore developed a set of features that reflect this basis. We used hexagonal lattices with spacings of 480 m, 240 m, 120 m, 80 m, 40 m, 20, and 10 m in physical space. For each lattice, the lattice was constructed by placing 2D Gaussian functions at the vertices. These Gaussian functions had a standard deviation of 1/12 the spacing of the lattice to account for the phase-offset copies of the lattice. Each lattice forms a basis function. For each spacing, we included variants with spatial phase offsets of ¼, ½, and ¾ the spacing, and also rotated copies at 20 and 40 degrees of rotation. The value of each basis function at each frame was computed by taking the maximum value of these associated 2D gaussian functions evaluated at the position of the participant. This feature space contains a total of 336 features that tiled the map.

*Head direction*

Head direction selectivity has been demonstrated in rodents [37] , non-human primates [38], and humans [17]. In this experiment, we defined the head direction as the orientation of the participant's car in the virtual world. At every frame, this orientation was then binned into one of eight equally spaced orientation bins (N, S, E, W, NE, NW, SE, SW) for a total of 8 features.

*6-fold head direction*

While grid cells are well-established in electrophysiology, they remain more elusive in fMRI. [18]demonstrated evidence for a grid-like representation indirectly using fMRI. This study is premised on the basis that the axes of individual grid cells are spatially aligned. Thus, travelling in a direction aligned with the hexagonal axes should cause grid cells to fire more than if the direction of travel is misaligned. Voxels containing grid cells should therefore display a six-fold symmetry in activity relative to head direction. [18]indeed shows such six-fold symmetry and thus suggests that it provides evidence for grid cells in fMRI.

We developed the 6-fold head direction feature space to capture this possible representation in our experiment. We constructed pairs of sinusoid functions with 90 degree phase offsets that sweep through 6 cycles across 360 degrees. Bins that are out of phase with the actual selectivity would perform poorly, whereas these functions would be able to capture the phase of the preferred direction. At every frame, we took the heading of the participant, and calculated the value of the two sinusoid functions at this direction for a total of 2 features.

*Gaze direction*

We defined the head direction as the allocentric heading of the participant's car in the virtual world. Because participants are not fixating at the center of the screen, they may not necessarily be looking in the direction of travel. The virtual camera has a horizontal field of view of 110°. Therefore, the allocentric gaze direction may differ from the car heading by up to 55°, a significant divergence. We also build features to account for a possibly separate representation of the allocentric orientation of the gaze direction in addition to the head direction. At every frame, this gaze direction was binned into one of eight orientation bins in the same manner as the head direction feature space, for a total of 8 features.

*6-fold gaze direction*

Similar to the difference between the gaze direction and head direction, there may also be separate six-fold symmetric representations for the gaze direction and head direction. Thus, at every frame, we computed the value of the same sinusoid functions used in the six-fold head direction feature space for the gaze direction of the participants, for a total of 2 features.

*Road graph*

Human behavioral studies suggest that larger spaces are represented on a relational basis that conserves their topologic organization, but not necessarily their metric spatial organization [19,20]. Furthermore, a neuroimaging study suggests that the brain represents graph-theoretic measures about road networks [21]. Thus, we built a graphical model of the road network in the virtual city. Intersections were used as vertices and road segments between intersections were used as edges for a total of 232 vertices and 294 edges. Furthermore, graphical planning may be a multiscale process, in which plans are conceptualized on both general (e.g., going to the grocery store) and specific (e.g., the particular route to take) levels. To capture this multiscale planning, we performed graph simplification on this road graph. In this process, the closest pair of vertices is merged into a single vertex, and the edge connecting the pair is deleted. All edges connecting to either of the two original vertices are replaced with new edges that connect to the new vertex. This step is iteratively applied to the graph until there remains only a single vertex. These new vertices and edges increasingly reflect the implicit connectivity between different parts of the map, rather than the direct road graph. Including the aggregate vertices and edges, there are a total of 463 vertices and 1042 edges. We use edges as features for a total of 1042 features.

At each frame, we determine the road edges that correspond to the path that the participant will take to the destination. We set the feature value corresponding to each of these edges to 1. These edges reflect the specific plan. Then, for the vertices that these edges connected, we find all aggregate vertices into which they collapse. We then find all edges that connect all these aggregate vertices, and also set their feature values to 1. These aggregate edge features reflect the more general plan. Taken together, these features reflect the participants' multiscale abstract navigation plan on the road network.

*Pedestrians & Other Vehicles*

Navigation in the real world is a dynamic process that may require interaction with, and therefore representation of, other agents. Indeed, electrophysiology studies show that both rat and bat hippocampus contain social place cells that represent the position of other animals in the environment [22,23]. It is likely that the human brain contains similar representations. In the virtual world used in this experiment, there are two types of AI entities – vehicles and pedestrians – both of which may influence the immediate navigational decisions of the participant. We developed a set of features to capture these possible representations.

At every frame, we used the demo files to extract the position and view vector of the participant, and also the position of all other vehicles and pedestrians in the world. Next, we constructed two identical sets of egocentric log-polar bins around the participant. For each of 16 directions, we placed 10 log-spaced distance bins from 5 m to 200 m away from the participant. For the first set of bins, we counted the number of vehicles in each bin. For the second set of bins, we counted the number of pedestrians in each bin. We separated the representation of vehicles and pedestrians into two feature spaces, because these two types of entities are visually distinct and behave differently. Thus, they may require distinct representations. Vehicle and pedestrian bins were treated as two separate feature spaces, each with 160 features.
